# Stalling of Transcription by Putative G-quadruplex Sequences and CRISPR-dCas9

**DOI:** 10.1101/2024.03.17.585391

**Authors:** Mohammed Enamul Hoque, Mohammad Lutful Kabir, Sajad Shiekh, Hamza Balci, Soumitra Basu

## Abstract

Putative G-quadruplex forming sequences (PQS) have been identified in promoter sequences of prominent genes that are implicated among others in cancer and neurological disorders. We explored mechanistic aspects of CRISPR-dCas9-mediated gene expression regulation, which is transient and sequence specific unlike alternative approaches that lack such specificity or create permanent mutations, using the PQS in tyrosine hydroxylase (*TH*) and *c-Myc* promoters as model systems. We performed *in vitro* ensemble and single molecule investigations to study whether G-quadruplex (GQ) structures or dCas9 impede T7 RNA polymerase (RNAP) elongation process and whether orientation of these factors is significant. Our results demonstrate that dCas9 is more likely to block RNAP progression when the non-template strand is targeted. While the GQ in *TH* promoter was effectively destabilized when the dCas9 target site partially overlapped with the PQS, the *c-Myc* GQ remained folded and stalled RNAP elongation. We also determined that a minimum separation between the transcription start site and the dCas9 target site is required for effective stalling of RNAP by dCas9. Our study provides significant insights about the factors that impact dCas9-mediated transcription regulation when dCas9 targets the vicinity of sequences that form secondary structures and provides practical guidelines for designing guide RNA sequences.

## INTRODUCTION

Clustered regularly interspaced short palindromic repeats (CRISPR) and the CRISPR associated (Cas) proteins are widely found in bacteria and archaea and serve as an RNA-mediated adaptive immune system that protects against invading microphages (1, 2). The use of an engineered nuclease-deficient Cas9, known as dCas9, allows the system to be repurposed for targeting genomic DNA without cleaving it (3). This repurposing was initially discovered by introducing mutations into the HNH and RuvC nuclease domains of the *S. pyogenes* Cas9 (3, 4). The resulting nuclease-deficient dCas9 and guide RNA (gRNA) complex can specifically bind to the target sequence and allows for direct manipulation of the transcription process without genetically affecting the DNA sequence. In order to activate and enhance the transcription of the target genes, the dCas9 fusion with a transcriptional activator domain can recruit RNA polymerase (RNAP) (5–8). The CRISPR-dCas9 mediated transcription activation (CRISPRa) is straightforward, target-specific, programmable, and frequently applicable when compared to conventional expression techniques that are based on the modification of genes and promoters (5, 9). It also enables the recruitment of various effector proteins for gene regulation at the transcriptional level (10, 11). In *E. coli*, CRISPR interference (CRISPRi) technology was used to show the effectiveness of dCas9 for sequence-specific gene repression. The dCas9-gRNA complex can interfere with transcription elongation by blocking RNAP (3, 12, 13). It can also obstruct transcription initiation by interfering with transcription factor binding (3, 4, 14, 15). Another benefit of using dCas9 is that it can direct its regulatory effect to a promoter instead of requiring the insertion of an operator sequence to control a native gene (16–18). It has been reported that targeting with multiple gRNAs in overlapping or different positions of a single promoter results in mutually exclusive binding that either recruits or blocks RNAP (13, 19–22). Although there are numerous *in cellulo* and *in vitro* examples of transcription regulation enacted by targeting one or more sites with dCas9-gRNA, to our knowledge there are no examples of mechanistic details how dCas9 behaves in the context of DNA secondary structures, such as the G-quadruplex (GQ).

The human genome contains an uneven distribution of putative GQ sequences (PQS) (23–25), and it has been shown that PQS located at promoter and 5′-UTR can regulate transcription and translation, respectively, *in vivo* (26). It is reported that the effect of PQS on transcriptional regulation is determined by several factors, including the sequence composition of the PQS, its location in relation to transcription start site, and its orientation (whether it is located in template or non-template strand) (27–29). Previously, we demonstrated that targeting the vicinity of a PQS in the human tyrosine hydroxylase (*TH*) promoter by dCas9 *in cellulo* causes transcription to be up or down regulated depending on the target sites (30). However, the mechanistic details of this process are not clear. The goal of this study is to investigate the interactions of RNAP and dCas9 in the vicinity of this PQS to gain mechanistic understanding of dCas9-mediated transcription regulation (Fig. 1A). To achieve this, we have designed different DNA and gRNA constructs, which were primarily used for *in vitro* transcription in an RNAP extension and single molecule Förster Resonance Energy Transfer (FRET) assays (31). In general, RNAP transcribes RNA until a full-length product is elongated (Fig. 1B). However, in the presence of PQS, RNAP interacts with the GQ structure and can be halted depending on the position and stability of the GQ (29, 32, 33), resulting in truncated RNA product rather than full-length RNA (Fig. 1C). In the presence of dCas9 targeting the vicinity of PQS, it may also serve as a blockade with or without destabilizing the GQ structure (Fig. 1D).

**Figure 1.**
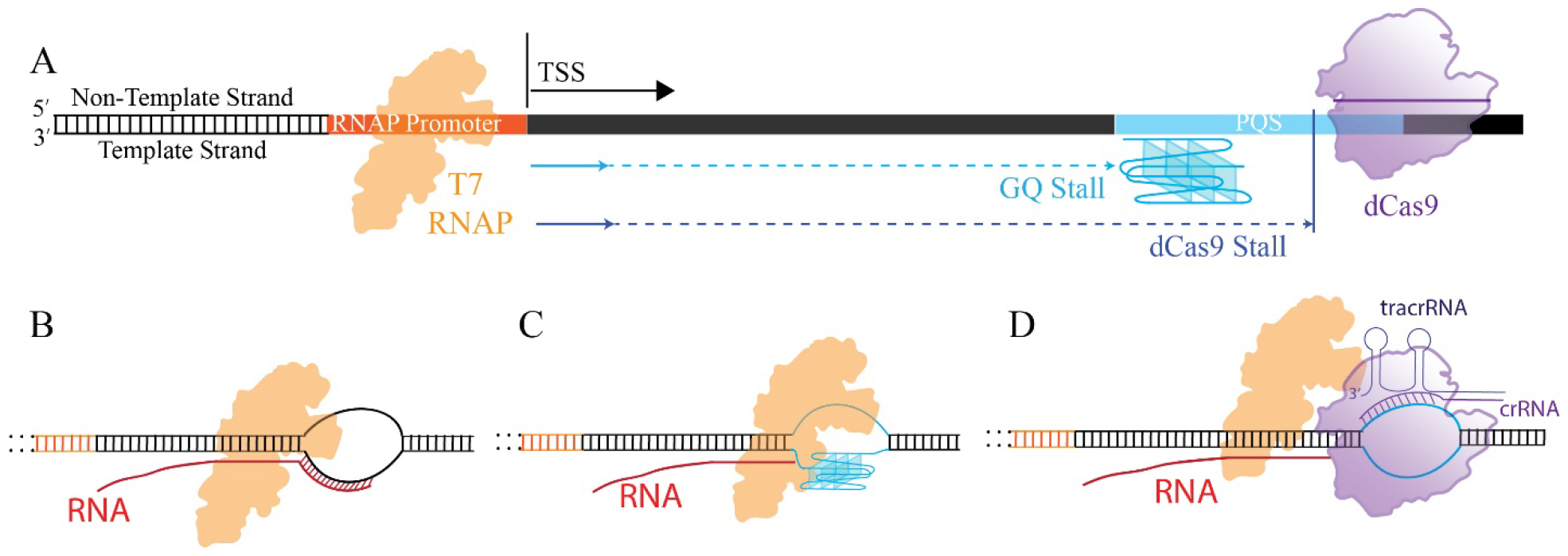
(A) Schematic of the DNA construct in which the impact of dCas9 and GQ on the transcription process was investigated. (B) In the absence of GQ and dCas9, the transcription is expected to proceed until a full-length RNA molecule is produced. (C) Formation of GQ in template strand is expected to present a blockade for RNAP, resulting in truncated RNA products of a specific length. (D) A schematic that demonstrates a case where dCas9 targets the non-template strand and presents a blockade for RNAP.

As model systems, we used the DNA constructs that contain the PQS from the TH (30, 34, 35), and the *c-Myc* (36) promoters. TH is the rate limiting enzyme in dopamine biosynthesis and is linked to various neurological disorders, including bipolar disorder (37) and schizophrenia (38, 39). C-Myc is an important oncoprotein and a transcription factor and plays an essential role in cell proliferation and induction of apoptosis (40–42). Overexpression of c-Myc is associated with a significant number of human malignancies, including breast, colon, cervix, and small-cell lung cancers, osteosarcomas, glioblastomas, and myeloid leukemias (43, 44). The c-Myc transcription is under the complex control of multiple promoters. The nuclease hypersensitivity element III1 (NHE III1) in the proximal region of the *c-Myc* promoter (−142 to −115 base pairs) can form two highly stable and competitive GQs controls 80–90% of the total transcriptional activity of this gene (45, 46). Similarly, the TH PQS contains five consecutive G-rich repeats, enabling formation of multiple GQs. In both systems, the PQS are placed downstream of the transcription start site (TSS) in order to quantify their potential inhibition of T7 RNAP based on truncated RNA products. For the TH system, we previously detected two prominent truncation sites for DNA polymerase that coincide with the GQ formation (35).

In the absence of a PQS, dCas9 blocks RNAP from *E. coli* and bacteriophages SP6, T3, and T7 to different extents (12). Even though both dCas9 and GQs can independently block RNAP progression, dCas9 alone may also stabilize or destabilize the GQs (depending on the target site) (47); thus, affecting transcription. Therefore, when PQS and dCas9 are present together, dCas9 may either promote RNAP progression by destabilizing a GQ structure or repress RNAP progression by further stabilizing the GQ structure or acting as a blockade by itself. Since the dCas9 targeting is reversible, these competing effects might enable dCas9 to up or down regulate transcription.

## METHODS

### Oligonucleotide preparation

All RNA and DNA oligonucleotides sequence information are reported in Tables S1, S2, and S3. The tyrosine hydroxylase PQS is: 5^′^-**<u>GGGG</u>**T**<u>GGGGG</u>**ATGTAA*GG*A **<u>GGGG</u>**AA*GG*T**<u>GGGGG</u>**ACCCAGA**<u>GGGGG</u>**, which contains five G-tracts identified with bold underlined letters. The GQ formed by the four consecutive G-tracts on the 5^′^ side (G-Tracts 1-4) is the most stable GQ followed by that formed by four consecutive G-tracts on the 3^′^ side (G-Tracts 2-5); although it is possible to form GQs with lower stability using other combinations of G-tracts (35). The sequence of the 27 nt long PQS of the c-Myc promoter is follows: T<u>GGGG</u>A<u>GGG</u>T<u>GGGG</u>A<u>GGG</u>T<u>GGGG</u>AAGG, with the underlined five G-tracts shown to form two different GQ structures (32).

We used separate strands for CRISPR-RNA (crRNA) and transactivating CRISPR RNA (tracrRNA) in the CRISPR-dCas9 complex. Annealing these two strands resulted in the guide RNA (gRNA). In case of TH, system, 5^′^ end amine modified or unmodified crRNA sequences crRNA oligos (crR-1, crR-2, crR-3, and crR-4) were purchased in 2^′^-protected form from Dharmacon, Inc. The 2^′^ protected RNAs were deprotected by using 2^′^-ACE deprotection buffer according to the manufacturer’s protocol. The tracrRNA and all crRNA components used for the c-Myc system were *in vitro* transcribed in the lab. All DNA oligonucleotides (including those used as template for *in vitro* transcription) were purchased from either Integrated DNA Technologies (IDT) or Eurofins Genomics. The DNA and RNA products were purified via denaturing polyacrylamide gel electrophoresis (PAGE) with different percentages. Full-length products were visualized by UV shadowing and were excised from the gel. The DNA and RNA were harvested via the crush and soak method by tumbling the gel slice overnight at 4 °C in a solution of 300 mM NaCl, 10 mM Tris-HCl, and 0.1 mM EDTA (pH 7.4). Salt was removed by ethanol precipitation of the oligonucleotides twice, with two cold 70% (v/v) ethanol washes in between each precipitation. The oligonucleotides were dissolved in nuclease free water and stored at -20 °C. The c-Myc constructs were prepared using the same protocols.

### *In vitro* transcription

TracrRNA, all of crRNA of c-Myc system, and some of crRNA oligonucleotides of TH system (crR-3T & crR-3C) were *in vitro* transcribed from T7 promoter containing template DNA by using T7 RNAP. 3 µM DNA template was transcribed in a 100 μL reaction in the presence of 1x transcription buffer (40 mM Tris-HCl, 2 mM spermidine, 10 mM DTT, and 6 mM MgCl_2_), 20 mM MgCl_2_, 2-6 mM NTPs (depending on the percentage of the individual nucleotides in the full-length transcribed RNA), and 10 μg/mL T7 RNAP. The transcribed RNAs were purified by loading them into different percentages of denaturing PAGE. UV shadowing was used to identify the full-length product band. The RNAs were extracted from the gel slice by soaking it overnight in elution buffer and then collecting the RNAs via ethanol precipitation, as previously described.

### Construct design when PQS is in the template strand of TH system

To investigate the competing effects of dCas9 and GQ structure during the *in vitro* transcription process, we designed two constructs (125 bp and 200 bp) for the TH system in which the PQS was kept in the template strand, which was chosen from TH promoter. The T7 promoter was then inserted into the 3^′^ side of the template strand and the 5^′^ side of the non-template strand. We used four crRNA sequences to target the vicinity of PQS with dCas9, which has a PAM sequence of 5^′^ NGG in the non-target strand. To form the 125 bp construct, we used 125 bp_TS (template strand) and 125 bp_NTS (non-template strand) sequences (see Fig. 2 for gRNA targeting sites).

**Figure 2.**
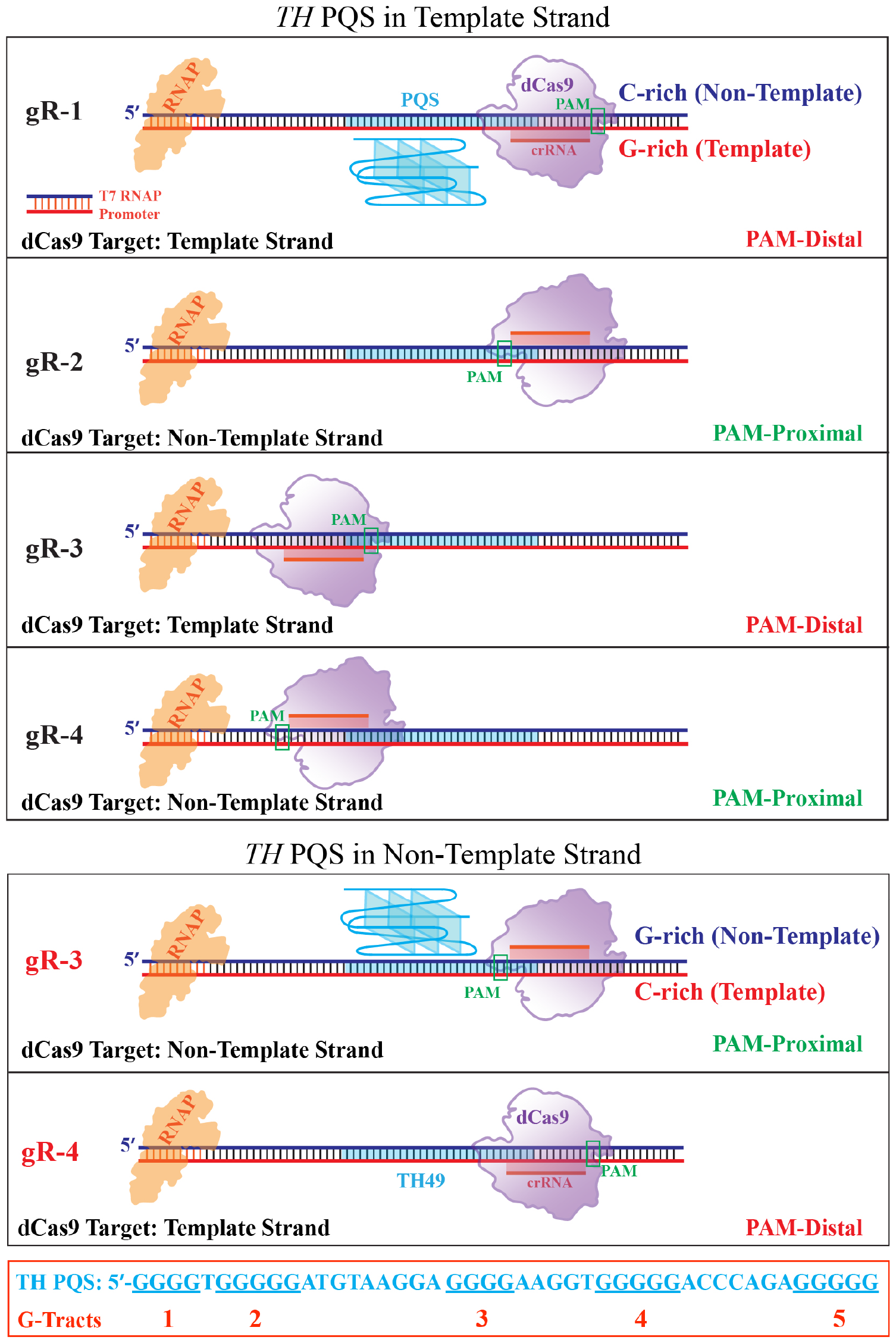
Schematics of the DNA constructs for the TH system when the putative quadruplex site is in the template (top) or non-template strand (bottom). Guide RNAs gR-1, gR-2, gR-3, and gR-4 are used in the former case while gR-3 and gR-4 are used for the latter. The bottom box provides the 45 nt long PQS located in TH promoter, which contains at least five G-tracts (of length 3 or more G’s) that can form multiple GQs.

To test whether the separation between RNAP promoter and dCas9 target site has any effect on RNAP progression, we inserted a sequence that moved the RNAP promoter 95 bps away from the first guide RNA binding site. We used 200 bp_TS (template strand) and 200 bp_NTS (non-template strand) to make this 200-bp construct.

In all these DNA constructs where PQS in the template strand, guide RNAs gR-1 and gR-3 were used as template strand targeting, while gR-2 and gR-4 were used as non-template strand targeting for CRISPR-dCas9 binding.

### Construct design when PQS is in the non-template strand of TH system

To evaluate the effect of GQ when it is located in the non-template strand, we designed a construct (135 bp) where the same PQS from *TH* promoter (same as that in the 125-bp and 200-bp constructs) is inserted in the non-template strand. To investigate *in vitro* transcription, the T7 RNAP promoter was inserted into the 3− side of the template strand and the 5^′^ side of the non-template strand. To use the same four guide RNAs (gR-1, gR-2, gR-3, and gR-4) to measure the impact of dCas9 on RNAP progression in this construct, the binding sites of these gRNA sequences were inserted accordingly. We used 135 bp_TS (template strand) and 135 bp_NTS (non-template strand) strands to build the 135-bp construct. In this construct, the guide RNAs gR-1 and gR-3 served as the non-template strand targeting, while gR-2 and gR-4 served as the template strand targeting for CRISPR-dCas9 binding.

### *In vitro* RNA polymerase assay

We used an *in vitro* T7 RNA polymerase assay to investigate the effect of GQ and dCas9 on transcription. The DNA construct was prepared by annealing the template strand and non-template strand in a 1:1 molar ratio (400 nM) at 95 °C for 5-10 minutes, followed by either slow or fast cooling to room temperature. The DNA strands were annealed under corresponding ionic conditions (K^+^ or Li^+^). The gRNA components crRNAs and tracrRNA were annealed separately (each at 1.2 µM) at 95 °C for 5 minutes followed by slow cooling to room temperature. The resulting guide RNA constructs are called gR-n accordingly. Ribonucleoprotein (RNP) complexes (dCas9-gRNA) were formed by mixing the annealed gRNA constructs with dCas9 protein (Sigma-Aldrich) in 2:1 molar ratio (600 nM) in the presence of 1xCas9 buffer (20 mM Tris-HCl, 100 mM KCl, 5 mM MgCl_2_, 1 mM DTT, and 5% glycerol).

The annealed DNA construct and the RNP complex were mixed and incubated at 37 °C for 20 minutes. Following incubation, *in vitro* transcription was performed by adding all of the transcription components. During *in vitro* transcription, 2-3 aliquots were removed at different reaction times and the transcription was terminated with stop buffer [7 M urea, 10 mM Tris-HCl, and 0.1 mM EDTA (pH 7.5)]. The reaction products were separated using 8% denaturing PAGE. The gel was stained with ethidium bromide solution for 25 minutes followed by visualization on a Typhoon FLA 9500 fluorescence imager (GE Life Sciences) by selecting Cy3 scanning mode. ImageJ software was used to further process the gel image.

### Single molecule experiments

Single molecule FRET measurements were performed using protocols described in detail in earlier work (48). PAGE or HPLC purified DNA oligos were purchased from Integrated DNA Technology (IDT). The DNA constructs consist of three stands including an 18 nt stem strand with biotin, another strand which includes the complementary sequence to the stem strand and an overhang, and a third strand that is complementary to this overhang. The DNA samples were annealed at 95 ºC for 5 minutes followed by slow cooling to room temperature.

A home-built prism type total internal reflection fluorescence microscope was used for single molecule experiments. The slides and coverslips were treated with 1M KOH followed by piranha etching. The slide surfaces were functionalized with amino silane solution followed by surface passivation with polyethylene glycol (PEG) over night. Additionally, the surfaces were treated with a second round of 333 kDa PEG to increase the density of the PEG brush and enhance the surface quality. Finally, the microfluidic chamber was created between a slide and cover slip with double-sided tape separating them. The chamber was washed with T50 buffer (10 mM Tris, 2 mM MgCl_2_ and 50 mM KCl or LiCl) before adding streptavidin followed by addition of the biotinylated DNA samples.

The DNA samples were diluted to 10-50 pM before adding to chamber to get the desired density of spots on the surface. Proteins and NTPs were also diluted to desired concentrations before they were introduced into the chamber. The imaging was done with a custom-built C++ program using an Andor Ixon EMCCD camera. We recorded short (15 frames) and long movies (1500 frames) in presence or absence of proteins and NTPs. The acquisition rate was at 100ms per frame. The data was analyzed in MATLAB, plotted and fitted in Origin.

## RESULTS

In the case of the TH system, which will be presented first, we designed several DNA constructs where the dCas9 target site and location of the PQS were varied. The PQS was placed in either the template or non-template strand for T7 RNAP and four guide RNA constructs (gR-1, gR-2, gR-3, and gR-4) were designed to target specific sites (Fig. 2). In the case of PQS in template strand, two DNA constructs, 125 bp (short) and 200 bp (long) in length, were utilized to test whether the separation between the RNAP and dCas9 binding sites influences the *in vitro* transcription process. For the short DNA construct, we also tested competitive binding to template and non-template strands. In the case of PQS in non-template strand, a 135-bp DNA construct was utilized. These measurements are followed with those on the c-Myc system in which the PQS was kept in the template strand as is the case in the physiological setting.

### I. The TH PQS System

In order to study the structures that are formed by the PQS before and after RNAP binding, and during RNAP progression, we conducted single molecule Förster resonance energy transfer (smFRET) experiments. CRISPR-dCas9 complex was not included in these studies to establish the baseline characteristics. The locations of the donor and acceptor fluorophores were optimized to be sensitive to structural changes around the PQS (Fig. 3A and Table S1). To establish a reference FRET level for the state where GQ is not folded (PQS hybridizes with the C-rich strand to form an intact dsDNA), we performed smFRET measurements at 150 mM LiCl, which does not stabilize the GQ structure. These data showed a single peak at FRET efficiency E_FRET_=0.50 (Fig. 3B). We then repeated these measurements in 50 mM KCl and observed two FRET peaks at E_FRET_=0.72 and E_FRET_=0.90 (Fig. 3B), which were both significantly higher than the reference peak in LiCl. We attributed these peaks to folding of (at least) two prominent GQs that were previously identified (35) and will be referred to as 5^′^-GQ or 3−-GQ depending on whether they contain the 1^st^-4^th^ G-tracts or the 2^nd^-5^th^ G-tracts of the PQS (counting from 5− side), respectively (see Table S1 for sequence). As there is minimal overlap between the peaks in LiCl and KCl conditions, we conclude that almost all DNA molecules contain folded GQ structures in KCl, before RNAP is added. Interestingly, RNAP binding increases the population of one of the GQs (the one represented with E_FRET_=0.90) while it decreases that of the other (Fig. 3C). Considering the proximity of the 5^th^ G-tract to the binding site of RNAP, the 3−-GQ is likely the destabilized structure. This suggests the disturbance created by RNAP binding is restricted to the 5^th^ G-tract and destabilization of the competing 3−-GQ facilitates folding of 5−-GQ. We then introduced NTPs to initiate the transcription elongation process (Fig. 3C). This resulted in emergence of a low FRET population (30% of total) that is consistent with the state observed in LiCl, i.e. the dsDNA where GQ is unfolded. The FRET peak identified as 5−-GQ largely remains intact and forms 70% of the total population. This suggests that about 30% of GQs are unfolded by T7 RNAP progression under these assay conditions while the others remain intact and could pose as blockade for T7 RNAP.

**Figure 3.**
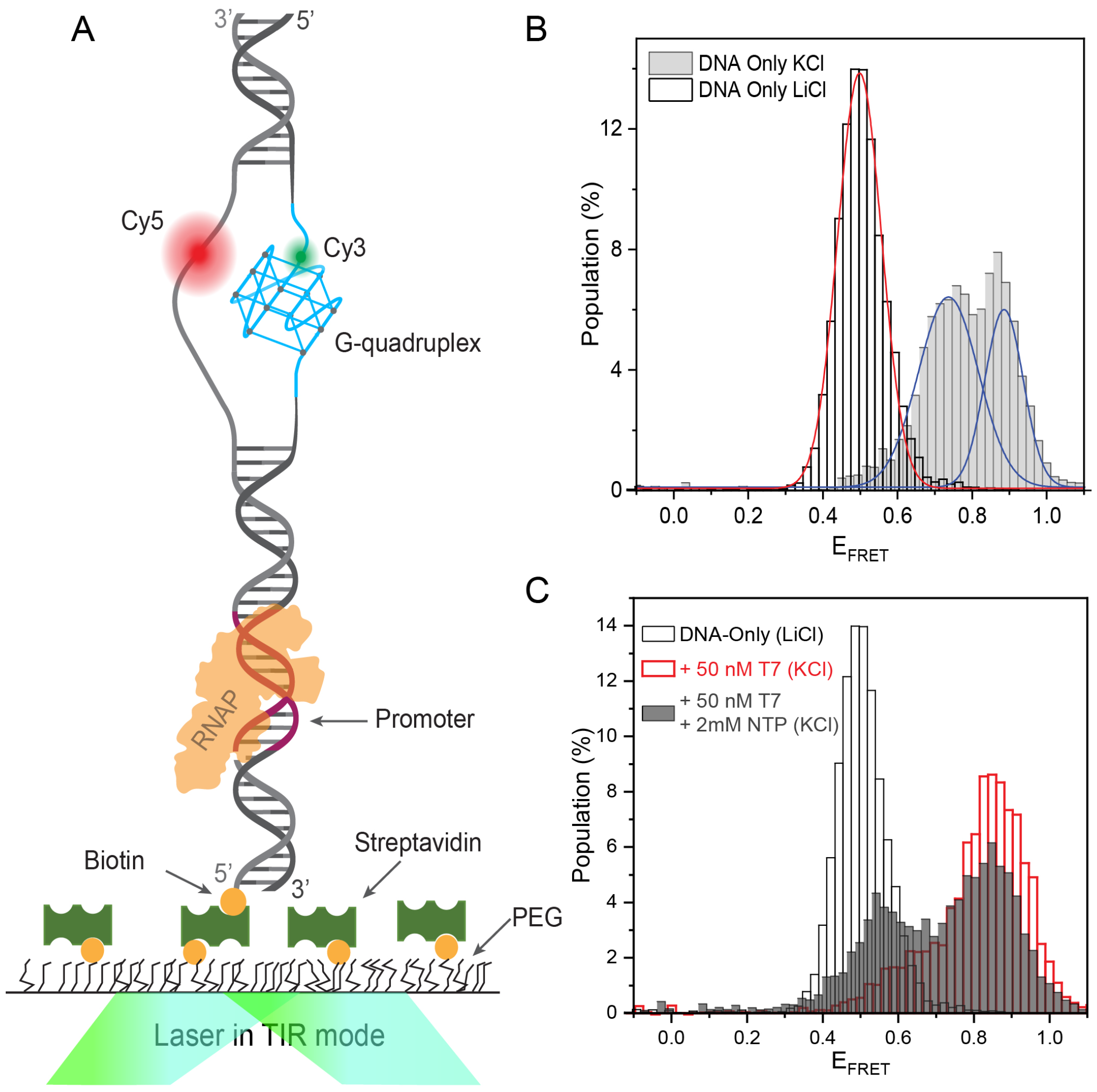
Single molecule FRET measurements demonstrating folding of GQ in TH system and activity of RNAP. (A) Schematic of the DNA construct showing the relative positions of important components of the assay. (B) smFRET histograms showing FRET peaks for the unfolded GQ case (in LiCl) and folded GQ case (in KCl). (C) smFRET histograms showing the shift in histograms upon introducing RNAP and nucleotides into the sample chamber.

#### I.A. *TH* PQS in the template strand

We performed *in vitro* transcription and RNAP stop assays on a 125-bp long DNA construct which contained the PQS in the template strand (Fig. 2 & Table S2). As shown in Fig. 4, in the absence of CRISPR-dCas9 (w/out dCas9 band) we were able to detect multiple truncated RNA products along with the full-length RNA bands in the PAGE assay in the presence of KCl. The truncated RNA bands correspond to the positions of the G-tracts, thus suggesting that the PQS folded into multiple GQ structures which halted RNAP progression at different sites (the length of these truncated products is given in Fig. 4 caption). The T7 RNAP arrest by GQ structure in the presence of K^+^ ions is also supported by previous findings (49). To confirm the truncated bands were due to the presence of GQ structures, we repeated these measurements in 100 mM LiCl or in the absence of additional salt which resulted in elimination or significant reduction of such truncated products (Fig. S1). As KCl is a more efficient stabilizer of the GQ(50), the stall bands are expected to be more prominent in KCl compared to LiCl.

**Figure 4.**
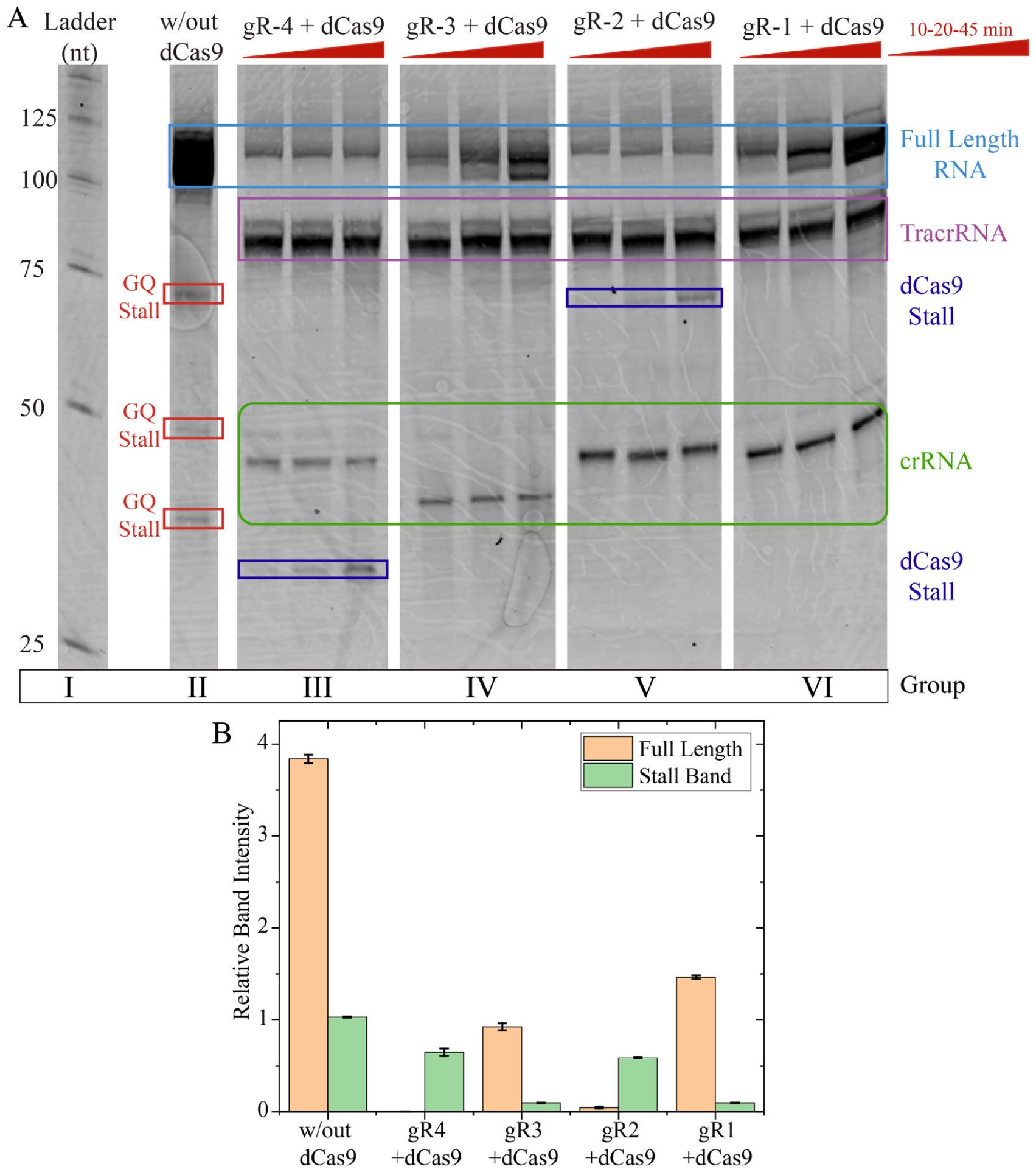
(A) Denaturing gel electrophoresis assay investigating the impact of dCas9 and PQS on RNAP progression when GQ is in the template strand for the TH case. A 125 bp construct was utilized in these experiments along with guide RNAs gR-1 & gR-3 (target template strand) and gR-2 & gR-4 (target the non-template strand). The truncated RNA products due to dCas9 stalls observed for gR-4 and gR-2 are expected to be around 26 nt and 80 nt long, respectively. The truncated RNA products due to GQ stalls in the absence of dCas9 are expected to be 39, 52, 60, and 73 nt long (depending on the position of the G-tracts and which GQ they form), three of which are detected and marked with red rectangles in Group II lane. The original (uncut) version of the gel is presented in Fig. S2. (B) Quantitation of full length and stall band intensities with respect to the template DNA band. In the case of GQ stalls in the absence of dCas9, all stall bands were combined for quantitation purposes. For gR-3 and gR1, in which there are no visible stall bands, the stall band intensity was determined by integrating the intensity at the expected stall location using a fixed-size rectangle.

To investigate the impact of dCas9 on RNAP elongation process, we performed *in vitro* transcription assays in the presence of CRISPR-dCas9 complexes. Guide RNA molecules gR-1 and gR-3 target the template strand, while gR-2 and gR-4 target the non-template strand (Fig. 2). The gR-1 was designed to block the first G-tract on the 5− side of the PQS from participation into the GQ structure, while gR-3 was designed to block the 3− side G-tract. Blocking a G-tract from taking part in GQ formation might facilitate GQ formation by the remaining G-tracts as it inhibits a competing structure or prevent formation of any GQ depending on the extent of the overlap. In the case of non-template targeting with gR-2 and gR-4, the R-loop formation between gRNA and the C-rich strand could unwind the dsDNA and facilitate folding of the GQs; although, how the presence of the CRISPR-dCas9 in close proximity of the GQ would impact unfolding of the GQ by RNAP is not known.

As illustrated in Fig. 4B, the transcription efficiency of the full-length RNA slightly decreased when the template strand was targeted with CRISPR-dCas9 (gR-1 and gR-3 cases in Fig. 4), which is consistent with reports on other systems (12, 13, 51); however, we did not detect any truncation products due to GQ or dCas9 stalls. In case of the non-template strand targeting by CRISPR-dCas9 (gR-2 and gR-4 cases in Fig. 4), we detected a significant reduction in the full-length RNA band and observed additional truncated bands due to dCas9 stalls (blue boxes) at the expected lengths, as quantified in Fig. 4B (band lengths given in Fig. 4 caption). To rule out the possibility of the direct effect of the guide RNA on the RNAP progression, we performed several transcription experiments in which gR-2 or gR-4 were included in the absence of dCas9. We did not detect the truncated bands in these studies (Fig. S2). These studies suggest dCas9 is more likely to block transcription when targeting the non-template strand (12, 13).

We did not observe a truncation band while targeting the template strand of the 125-bp DNA construct with dCas9; however, we observed a reduction in the full length product, in agreement with earlier research indicating weak repression (12). In this 125 bp DNA construct, the RNAP promoter is only 28 bps away from the dCas9 binding site, which raised concerns about whether this was sufficient separation for dCas9 to impact the transcription. Therefore, we designed a longer 200-bp DNA construct with the promoter 95 bps away from the first guide RNA binding site and repeated the *in vitro* transcription assays (Fig. 5). In the absence of CRISPR-dCas9, we detected multiple truncated RNA products, attributed to GQ stalls, along with the full-length RNA bands in the PAGE assay, which is consistent with the measurements on the 125-bp construct. We also observed a few truncated bands in the 200-bp construct that did not correspond to the positions of the G-tracts, implying the presence of misfolded structures, possibly induced by GQ formation, that could halt RNAP progression.

**Figure 5.**
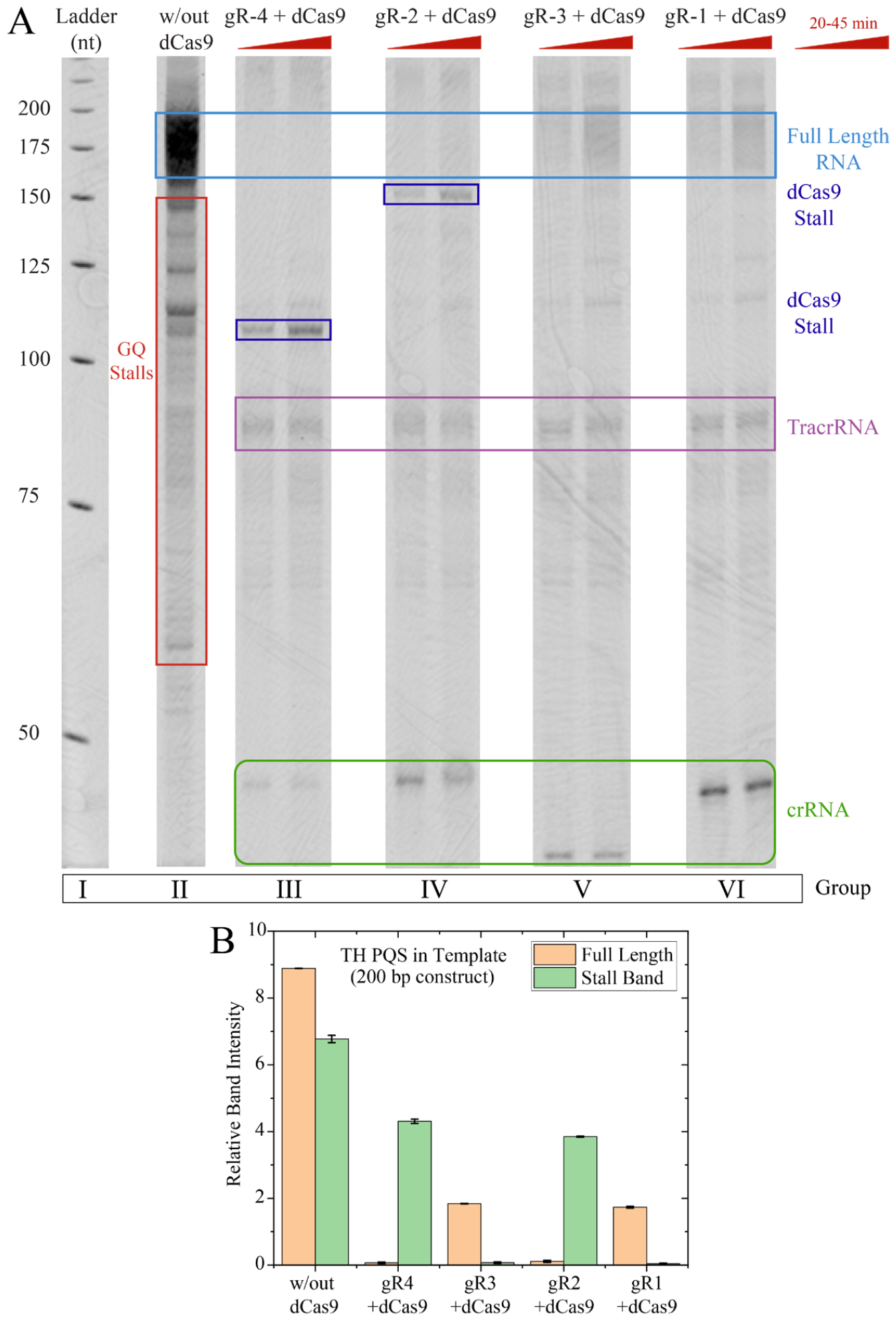
(A) Denaturing gel electrophoresis assay investigating the impact of dCas9 and PQS on RNAP progression when GQ is in the template strand for a 200 bp template DNA construct (longer length to investigate the effect of the separation between TSS and dCas9 target site on transcription). Guide RNAs gR-1 & gR-3 target the template strand and gR-2 & gR-4 the non-template strand. The truncated RNA products due to dCas9 stalls observed for gR-4 and gR-2 are expected to be around 95 nt and 149 nt long, respectively, which are consistent with the marked stalls in blue rectangles. The original (uncut) version of the gel is presented in Fig. S3. (B) Quantitation of full length and stall band intensities with respect to the template DNA band intensity. In the case of GQ stalls in the absence of dCas9, all stall bands were combined for quantitation purposes. For gR-3 and gR1, in which there are no visible stall bands, the stall band intensity was determined by integrating the intensity at the expected stall location using a fixed size rectangle. The error bars are based on standard deviation of at least three measurements.

We then targeted the vicinity of the GQ structure with CRISPR-dCas9 by using the same four guide RNAs (gR-1, gR-2, gR-3, and gR-4) and repeated the *in vitro* transcription assays (Fig. 5 and Fig. S3). In this case, the intensity of the full-length RNA product band significantly decreased when the template strand was targeted by gR-1 or gR-3 complexes (quantified in Fig. 5B), but we did not detect any additional stalls or truncation products. We also observed a significant reduction in full-length RNA band (barely visible) when the non-template strand was targeted with gR-2 or gR-4 complexes, and we detected truncated bands due to dCas9 blocks at the expected length (truncated product lengths given in Fig. 5 caption).

In the case of gR-4 targeting the 200-bp construct, the dCas9 stalls had significantly higher truncated band intensity (about two-fold higher when normalized with respect to corresponding template DNA band intensity) than that in the 125 bp construct (Fig. 5B vs. Fig. 4B). These studies suggest the requirement of a minimum separation between the RNAP promoter and dCas9 binding site for dCas9 to play a more prominent role in the elongation repression during *in vitro* transcription. Despite these interesting findings, more systematic studies are required to better understand the nature of this effect.

#### I.B. *TH* PQS in the non-template strand

We designed a 135-bp DNA construct in which the PQS is located in the non-template strand and performed similar *in vitro* transcription assays using gR-3 and gR-4 constructs (Fig. 6). In the absence of CRISPR-dCas9, we observed predominantly full-length RNA bands on the polyacrylamide gel. We did not detect any truncated product bands corresponding to the PQS regions, indicating that there were no GQ stalls when the PQS was in the non-template strand. It has been suggested that GQ structures located in the non-template strand promote transcription rather than hinder RNAP elongation(29), and our studies indicate that at least they do not hinder the elongation process. Then, we used gR-3 to target the non-template strand and gR-4 to target the template strand with dCas9 (Fig. 6, notice the switch in the orientation of gR-3 and gR-4 compared to the earlier case). In case of targeting the template strand, we did not detect any additional stalls or truncation products, but just the full-length RNA bands^13,14^. However, in case of targeting the non-template strand with gR-3, we found a significant reduction in the full-length RNA band and observed truncated bands (blue box) at the length expected for dCas9 stall. When dCas9 was eliminated from this reaction (gR-3 without dCas9), we did not detect any additional truncated bands, indicating that dCas9 blocks transcription when targeting the non-template strand (Fig. S4)(12, 13).

**Figure 6.**
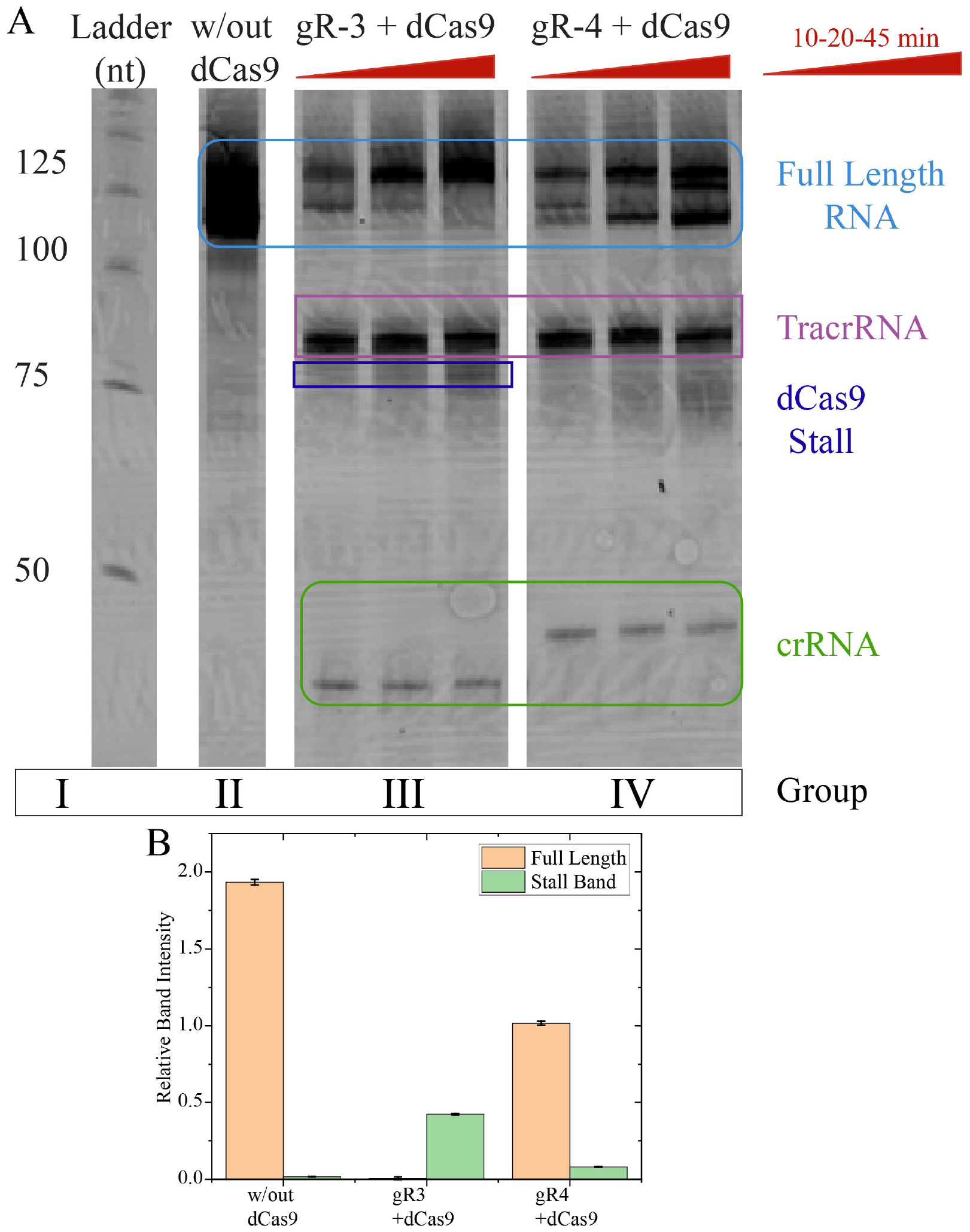
Denaturing gel electrophoresis assay investigating the impact of dCas9 and PQS on RNAP progression when TH PGQ is in the non-template strand. A 135 bp construct was utilized in these experiments along with the guide RNAs gR-3 and gR-4, which target the non-template and template strands, respectively (notice the switch in orientation compared to earlier constructs). The original (uncut) version of the gel is presented in Fig. S4. (B) Quantitation of full length and stall band intensities with respect to the template DNA band intensity. In cases where there were no visible stall bands, the stall band intensity was determined by integrating the intensity at the expected stall location using a fixed size rectangle. The error bars are based on standard deviation of at least three measurements.

#### I.C. Competitive binding of gRNAs between template and non-template strands

We also investigated the case of targeting both strands simultaneously using CRISPR-dCas9 (Fig. 7 and Fig. S5). To do so, we utilized gR-3 and gR-4 guide RNA constructs since their PAM sequences are only 5 bps apart. These measurements were performed on the TH PQS in template strand construct (the 125 bp long construct of Fig. 4). We first incubated the reaction mixture with [gR-3+dCas9] complex for 15 minutes before adding gR-4 RNA only (without dCas9) ([gR-3+dCas9] + gR-4), followed by aliquoting the reaction mixtures at three different time points. The goal of these measurements were to investigate whether dCas9 would dissociate from gR-3 (which targets the template strand) and bind to gR-4 (which targets the non-template strand). We observed a reduction in the full-length RNA bands and the appearance of the dCas9 stalls at the same location as that observed for gR-4 case (Fig. 4), which were not present in the case of gR-3. This observation indicates that the dCas9 switched from template strand targeting site (gR-3) to non-template targeting site (gR-4). We then reversed the order by incubating the reaction mixture with [gR-4+dCas9] complex for 15 minutes before adding gR-3 RNA ([gR-4/dCas9] + gR-3) and aliquoted the reaction mixtures similarly. In this case, adding gR-3 RNA later did not appear to impact the results and the dCas9 blocks were consistent with those observed for gR-4 case. Therefore, the non-template strand targeting guide RNA (gR-4) appears to be more dominant over template strand targeting (gR-3) during *in vitro* transcription.

**Figure 7.**
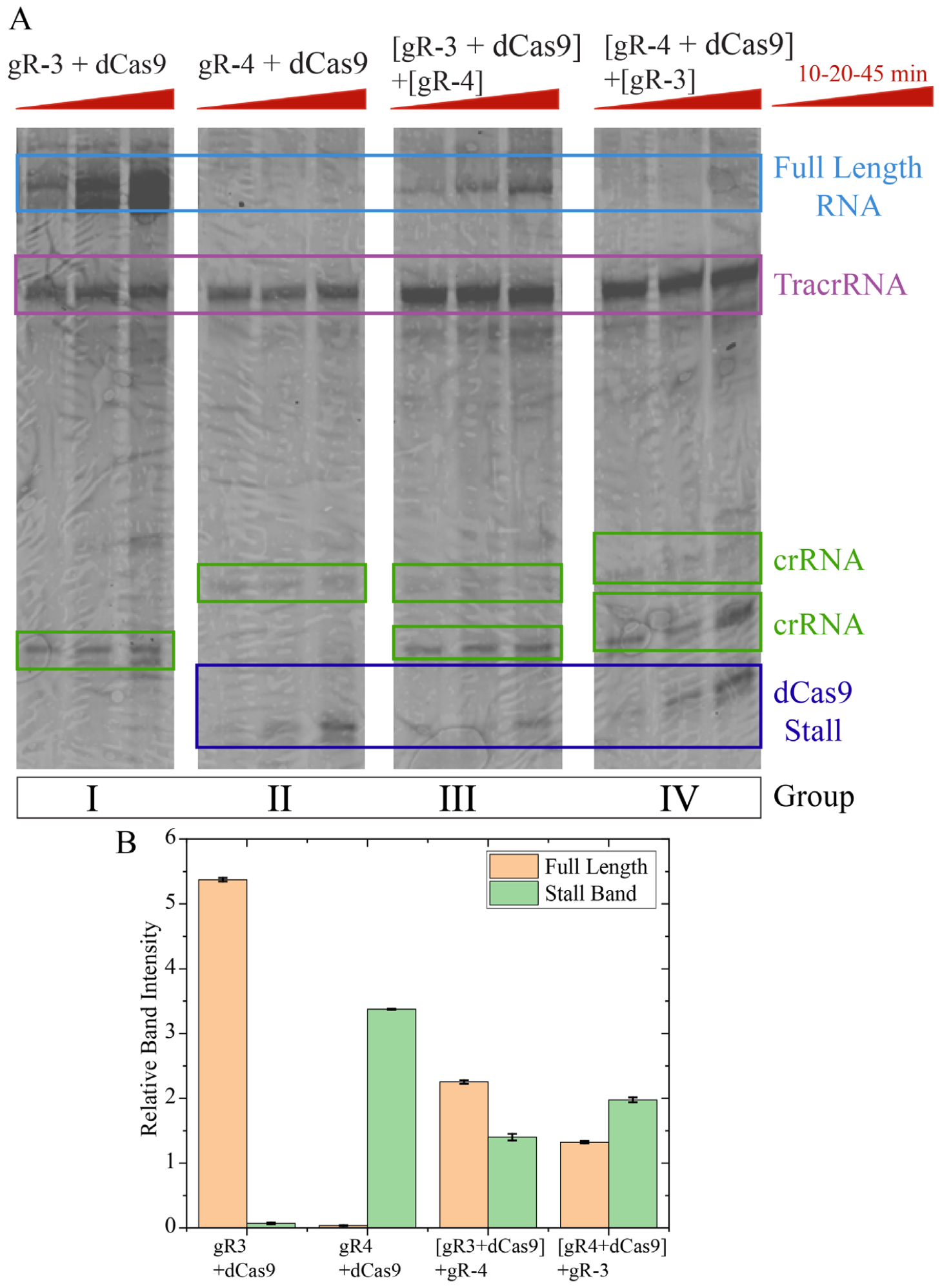
Competitive binding of guide RNAs to the template vs. non-template strands. The TH PQS in template strand construct (that of Fig. 4) was used in these measurements. Complete complexes (gRNA+dCas9) or just the guide RNAs (gR-3 & gR-4) are targeted to the opposite strands during RNAP progression to investigate whether the dCas9 dissociates from its guide RNA (complete complex) and binds to the other gRNA which is not bound to a dCas9. When both guide RNA’s are incorporated into the assay, regardless of in which order they are introduced, the resulting pattern is consistent with that of gR-4+dCas9 case, which targets the non-template strand. The original (uncut) version of the gel is presented in Fig. S5. (B) Quantitation of full length and stall band intensities with respect to the template DNA band intensity. In cases where there were no visible stall bands, the stall band intensity was determined by integrating the intensity at the expected stall location using a fixed size rectangle. The error bars are based on standard deviation of at least three measurements.

### II. The *c-Myc* PQS System

We created a 194-bp long DNA construct that includes the T7 RNAP promoter, the 27-nt long c-Myc PQS (which contains five G-tracts) in the template strand, and the flanking sequences around the PQS. GQ formation in this construct was confirmed with circular dichroism measurements (Fig. S6). Taking the physiological sequence as the reference, we identified seven sites that can be targeted by CRISPR-dCas9 in the template and non-template strands in this construct (Fig. 8A). Some of these sites overlapped with the PQS or the complementary C-rich strand while others were upstream and downstream of it (without overlap). Specifically, in construct 1, we targeted the template strand such that the gR-1 was complementary to 1^st^ and 2^nd^ G-repeats of the PQS. In the 2^nd^ and 3^rd^ constructs, we targeted the non-template strand and gR-2 and gR-3 overlapped with sequences that were complementary to the G-repeats in the 5− and 3− sides, respectively. The other four constructs were designed to target the C-rich or the G-rich strands at the 5− and 3− sides that are away from the PQS.

**Figure 8.**
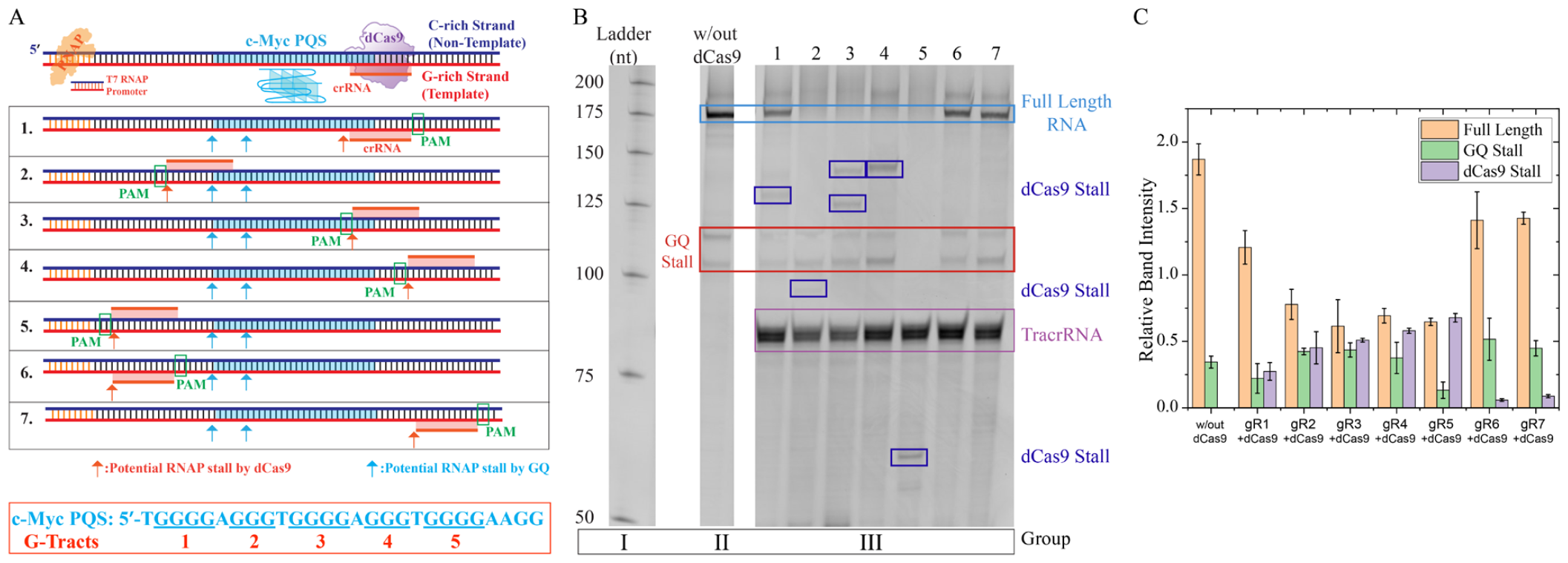
(A) Schematic of the DNA construct and seven different sites targeted by CRISPR-dCas9 in the c-Myc system. The PQS of the c-Myc system is given at the bottom box. (B) Denaturing gel electrophoresis assay investigating the impact of GQ structure and CRISPR-dCas9 complexes on transcription in the c-Myc system. For brevity gRNA and dCas9 complexes are marked with the corresponding numbers 1-7, which are described in (A). The RNAP stalls mediated by GQ and dCas9 are shown in red and blue boxes, respectively. The expected size of the truncated transcription products are as follows: GQ Stalls:106 nt and 11 nt (in all cases except Case 5); dCas9 stall in Case 1: 123 nt; dCas9 stall in Case 2: 102 nt; dCas9 stall in Case 3: 128 nt; dCas9 stall in Case 4: 149 nt; dCas9 stall in Case 5: 66 nt; dCas9 stall in Case 6: 68 nt; and dCas9 stall in Case 7: 144 nt. (C) Quantitation of the full length and stall band intensities with respect to the template DNA band intensity. The two GQ stall bands were combined for quantitation purposes. The error bars are based on standard deviation of at least three measurements.

The results of RNAP stop assay on the c-Myc system are shown in Fig. 8, where the GQ and dCas9 stall sites are indicated on the gel (Fig. 8B), quantified in Fig. 8C, and the expected lengths of the truncated RNA products are given in the caption. We observed stalls corresponding to the formation of two GQs (appear as two bands in the gel) in all the cases except for gR-5. In case of gR-5, the polymerase encounters and is stalled by the dCas9, which is bound to the non-template strand, before it reaches the GQ. In agreement with the TH case, we observed dCas9 stalls in all four cases (gR-2, gR-3, gR-4, and gR-5) when dCas9 targeted the non-template strand. On the other hand, we observed dCas9 stall in only one of the three cases (gR-1) when dCas9 targeted the template strand. Differently from the TH case, the GQ stalls were not eliminated when the vicinity of PQS was targeted by dCas9.

## DISCUSSION

In this study, we report a mechanistic approach to investigate how dCas9 impacts *in vitro* transcription in the presence of a GQ structure. To systematically study this question, we used model PQS systems from *TH* and *c-Myc* and targeted their vicinity with CRISPR-dCas9. In the case of TH, the PQS was placed in the template or non-template strands while the *c-Myc* PQS was kept in the template strand, as it is the case physiologically.

In the TH system and in the absence of dCas9, the GQ structure did not inhibit RNAP progression when the PQS was in the non-template strand (29); however, we observed several RNA truncation bands when PQS was in the template strand, suggesting PQS folded into multiple GQs which stalled RNAP progression (we have previously reported that the TH PQS can fold into multiple GQs and our current data are fully consistent with that finding (35)). When this template was targeted by CRISPR-dCas9, transcription efficiency of the full-length RNA product was significantly reduced, and we did not detect any truncated product bands beyond very faint bands in the gel. We attributed the weakening of the GQ stall bands to destabilization of the GQ by dCas9. We also observed that the impact of dCas9 was more significant when the separation between RNAP promoter and dCas9 binding site increased from 28 bp to 95 bp (DNA construct length increased from 125 bp to 200 bp). When the non-template strand is targeted with CRISPR-dCas9, we found a termination of RNAP elongation at sites corresponding to dCas9 binding, as well as significant reduction in full-length RNA product bands.

In the case of *c-Myc*, in which the PQS was in the template strand, we observed GQ stall bands in almost all cases we investigated, even when dCas9 targeted the proximity of these sites, which was different from that of the TH system. This difference could be due to the TH GQ being destabilized to a greater extent by the dCas9 and the R-loop between the gRNA and the G-rich DNA template. Consistently with the TH system, we observed dCas9 stalls in all cases when the dCas9 targeted the non-template strand. We observed such a stall in only one of three cases (and at a much lower intensity as quantified in Fig. 8C) when dCas9 targeted the template strand. Therefore, weak stalling of RNAP when dCas9 targets the template strand should be considered as a possibility.

An important question in this context is how dCas9 plays differential roles while targeting template or non-template strands (52, 53). As illustrated in the schematic in Fig. 2, there are significant differences between the two cases in the way dCas9 and RNAP interact with each other. When RNAP approaches the CRISPR-dCas9 complex that is bound to the template strand, the contact between RNAP and dCas9 happens on the PAM-distal side (PAM is located on the opposite side of the first interaction site as shown in Fig. 2). In this case, dCas9 complex appears to be dislodged from the template strand and T7 RNAP completes the transcription process (13). As a result, we observe the full-length product but with a reduced yield, possibly because of the delay in dislodging the dCas9 complex. On the other hand, when dCas9 binds to the non-template strand (Fig. 2), contact between RNAP and dCas9 takes place on PAM proximal site (PAM acting as the first interaction site). In this case, progression of T7 RNAP is more likely to be halted and RNAP is dislodged from the DNA, which results in truncated products (12, 13, 51). These ideas are summarized in the schematic in Fig. 9.

**Figure 9.**
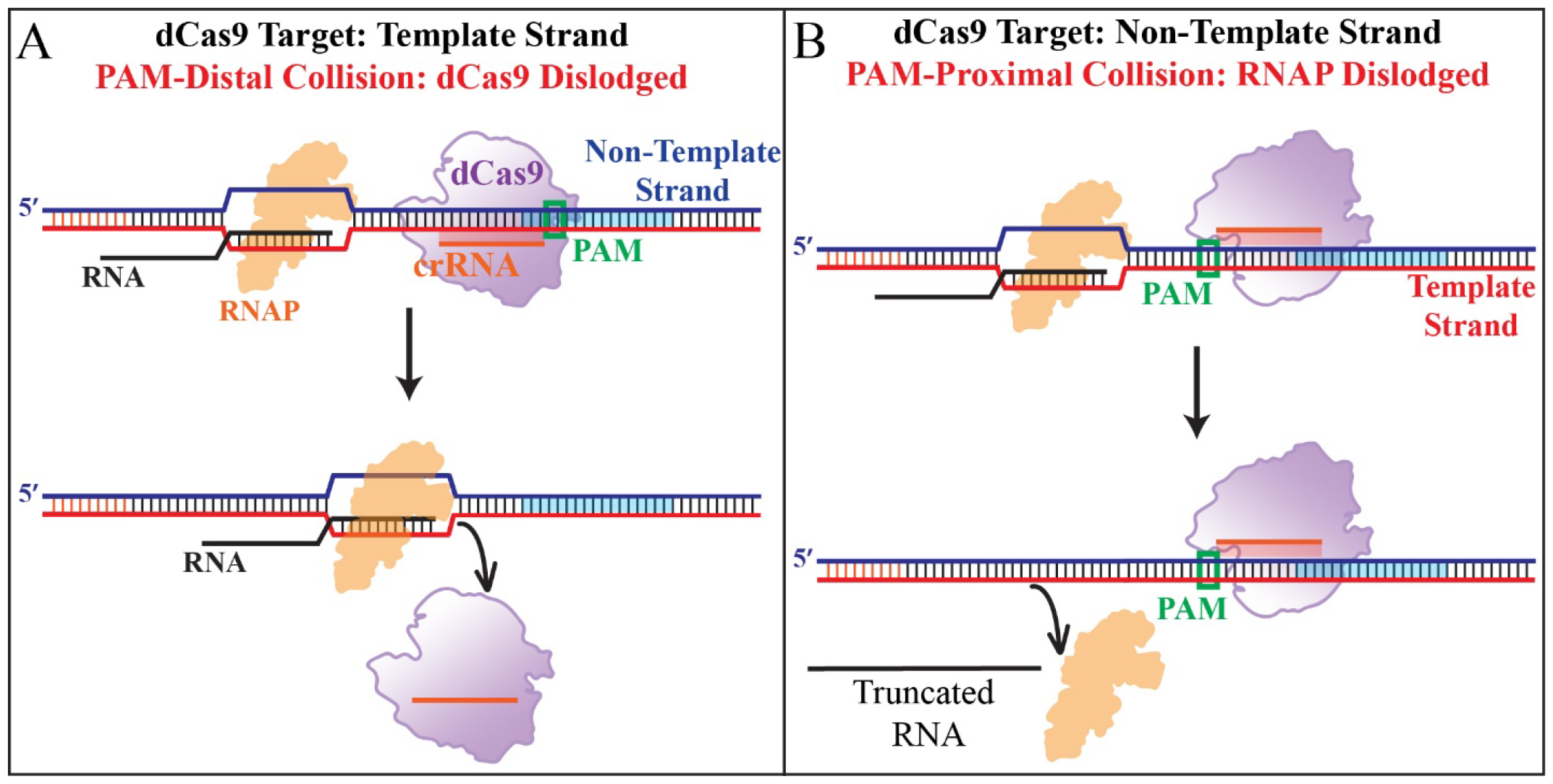
The mechanisms of interactions between dCas9 and RNAP in the vicinity of a PQS: A) When dCas9 targets the template strand, the collision between RNAP and dCas9 takes place in PAM-distal manner, which is more likely to result in dislodging of dCas9. (B) When dCas9 targets the non-template strand, the collision between RNAP and dCas9 takes place in PAM-proximal manner, which is more likely to result in dislodging of RNAP and truncated RNA products.

Our study provides important mechanistic insights about utilizing the CRISPR-dCas9 system for controlling gene expression by targeting the putative quadruplex sequences in two systems of medical significance. This technology can be used to specifically target and modulate gene expression, particularly in cases where GQs are known to form in the template strand and potentially interfere with transcriptional activity. When PQS is in the template strand, our data show that targeting the non-template strand with CRISPR-dCas9 results in dCas9 to be the more dominant block (compared to GQ) for RNAP progression. When GQ is in the template strand and this strand is targeted by CRISPR-dCas9, different levels of GQ destabilization are observed in TH and c-Myc systems. While the GQ blockade was almost completely eliminated by dCas9 in *TH*, it caused a significant stall in case of *c-Myc*. The ability to control gene expression in a site-specific manner has implications for a variety of research areas, including synthetic biology, gene therapy, and drug discovery. Our study highlights the need for further studies to comprehensively explore the exciting possibility of transiently controlling gene expression through targeting putative quadruplex sequences by CRISPR-dCas9 in a sequence specific manner.

## SUPPLEMENTARY DATA

Supplementary Data are available at NAR online.

## References

1. Horvath, P. and Barrangou, R. (2010) CRISPR/Cas, the Immune System of Bacteria and Archaea. Science (1979), 327, 167–170.

2. Barrangou, R., Fremaux, C., Deveau, H., Richards, M., Boyaval, P., Moineau, S., Romero, D.A. and Horvath, P. (2007) CRISPR Provides Acquired Resistance Against Viruses in Prokaryotes. Science (1979), 315, 1709–1712.

3. Qi, L.S., Larson, M.H., Gilbert, L.A., Doudna, J.A., Weissman, J.S., Arkin, A.P. and Lim, W.A. (2013) Repurposing CRISPR as an RNA-Guided Platform for Sequence-Specific Control of Gene Expression. Cell, 152, 1173–1183.

4. Larson, M.H., Gilbert, L.A., Wang, X., Lim, W.A., Weissman, J.S. and Qi, L.S. (2013) CRISPR interference (CRISPRi) for sequence-specific control of gene expression. Nat Protoc, 8, 2180–2196.

5. Konermann, S., Brigham, M.D., Trevino, A.E., Joung, J., Abudayyeh, O.O., Barcena, C., Hsu, P.D., Habib, N., Gootenberg, J.S., Nishimasu, H., et al. (2015) Genome-scale transcriptional activation by an engineered CRISPR-Cas9 complex. Nature, 517, 583–588.

6. Bikard, D., Jiang, W., Samai, P., Hochschild, A., Zhang, F. and Marraffini, L.A. (2013) Programmable repression and activation of bacterial gene expression using an engineered CRISPR-Cas system. Nucleic Acids Res, 41, 7429–7437.

7. Gilbert, L.A., Horlbeck, M.A., Adamson, B., Villalta, J.E., Chen, Y., Whitehead, E.H., Guimaraes, C., Panning, B., Ploegh, H.L., Bassik, M.C., et al. (2014) Genome-Scale CRISPR-Mediated Control of Gene Repression and Activation. Cell, 159, 647–661.

8. Chavez, A., Scheiman, J., Vora, S., Pruitt, B.W., Tuttle, M. P R Iyer, E., Lin, S., Kiani, S., Guzman, C.D., Wiegand, D.J., et al. (2015) Highly efficient Cas9-mediated transcriptional programming. Nat Methods, 12, 326–328.

9. Chen, S., Sanjana, N.E., Zheng, K., Shalem, O., Lee, K., Shi, X., Scott, D.A., Song, J., Pan, J.Q., Weissleder, R., et al. (2015) Genome-wide CRISPR Screen in a Mouse Model of Tumor Growth and Metastasis. Cell, 160, 1246–1260.

10. Dominguez, A.A., Lim, W.A. and Qi, L.S. (2016) Beyond editing: repurposing CRISPR–Cas9 for precision genome regulation and interrogation. Nat Rev Mol Cell Biol, 17, 5–15.

11. He, M., Zhou, X., Li, Z., Yin, X., Han, W., Zhou, J., Sun, X., Liu, X., Yao, D. and Liang, H. (2022) Programmable Transcriptional Modulation with a Structured RNA-Mediated CRISPR-dCas9 Complex. J Am Chem Soc, 144, 12690–12697.

12. Widom, J.R., Rai, V., Rohlman, C.E. and Walter, N.G. (2019) Versatile transcription control based on reversible dCas9 binding. RNA, 25, 1457–1469.

13. Anderson, D.A. and Voigt, C.A. (2021) Competitive dCas9 binding as a mechanism for transcriptional control. Mol Syst Biol, 17, e10512.

14. Bikard, D., Jiang, W., Samai, P., Hochschild, A., Zhang, F. and Marraffini, L.A. (2013) Programmable repression and acvtivation of bacterial gene expression using an engineered CRISPR-Cas system. Nucleic Acids Res, 41, 7429–7437.

15. Ji, W., Lee, D., Wong, E., Dadlani, P., Dinh, D., Huang, V., Kearns, K., Teng, S., Chen, S., Haliburton, J., et al. (2014) Specific gene repression by CRISPRi system transferred through bacterial conjugation. ACS Synth Biol, 3, 929–931.

16. Komor, A.C., Kim, Y.B., Packer, M.S., Zuris, J.A. and Liu, D.R. (2016) Programmable editing of a target base in genomic DNA without double-stranded DNA cleavage. Nature, 533, 420–424.

17. Tian, T., Kang, J.W., Kang, A. and Lee, T.S. (2019) Redirecting Metabolic Flux via Combinatorial Multiplex CRISPRi-Mediated Repression for Isopentenol Production in Escherichia coli. ACS Synth Biol, 8, 391–402.

18. Wu, Y., Chen, T., Liu, Y., Tian, R., Lv, X., Li, J., Du, G., Chen, J., Ledesma-Amaro, R. and Liu, L. (2020) Design of a programmable biosensor-CRISPRi genetic circuits for dynamic and autonomous dual-control of metabolic flux in Bacillus subtilis. Nucleic Acids Res, 48, 996–1009.

19. Qi, L.S., Larson, M.H., Gilbert, L.A., Doudna, J.A., Weissman, J.S., Arkin, A.P. and Lim, W.A. (2013) Repurposing CRISPR as an RNA-Guided Platform for Sequence-Specific Control of Gene Expression. Cell, 152, 1173–1183.

20. Lebar, T. and Jerala, R. (2016) Benchmarking of TALE- and CRISPR/dCas9-Based Transcriptional Regulators in Mammalian Cells for the Construction of Synthetic Genetic Circuits. ACS Synth Biol, 5, 1050–1058.

21. Gander, M.W., Vrana, J.D., Voje, W.E., Carothers, J.M. and Klavins, E. (2017) Digital logic circuits in yeast with CRISPR-dCas9 NOR gates. Nat Commun, 8, 15459.

22. Wu, F., Shim, J., Gong, T. and Tan, C. (2020) Orthogonal tuning of gene expression noise using CRISPR-Cas. Nucleic Acids Res, 48, e76.

23. Huppert, J.L. and Balasubramanian, S. (2007) G-quadruplexes in promoters throughout the human genome. Nucleic Acids Res, 35, 406–413.

24. Chambers, V.S., Marsico, G., Boutell, J.M., Di Antonio, M., Smith, G.P. and Balasubramanian, S. (2015) High-throughput sequencing of DNA G-quadruplex structures in the human genome. Nat Biotechnol, 33, 877–881.

25. Hänsel-Hertsch, R., Beraldi, D., Lensing, S. V, Marsico, G., Zyner, K., Parry, A., Di Antonio, M., Pike, J., Kimura, H., Narita, M., et al. (2016) G-quadruplex structures mark human regulatory chromatin. Nat Genet, 48, 1267–1272.

26. Halder, K., Wieland, M. and Hartig, J.S. (2009) Predictable suppression of gene expression by 5 ′-UTR-based RNA quadruplexes. Nucleic Acids Res, 37, 6811–6817.

27. Rigo, R., Palumbo, M. and Sissi, C. (2017) G-quadruplexes in human promoters: A challenge for therapeutic applications. Biochim Biophys Acta Gen Subj, 1861, 1399–1413.

28. Onel, B., Carver, M., Wu, G., Timonina, D., Kalarn, S., Larriva, M. and Yang, D. (2016) A New G-Quadruplex with Hairpin Loop Immediately Upstream of the Human BCL2 P1 Promoter Modulates Transcription. J Am Chem Soc, 138, 2563–2570.

29. Lee, C.-Y., McNerney, C., Ma, K., Zhao, W., Wang, A. and Myong, S. (2020) R-loop induced G-quadruplex in non-template promotes transcription by successive R-loop formation. Nat Commun, 11, 3392.

30. Hoque, M.E., Mustafa, G., Basu, S. and Balci, H. (2021) Encounters between Cas9/dCas9 and G-Quadruplexes: Implications for Transcription Regulation and Cas9-Mediated DNA Cleavage. ACS Synth Biol, 10, 972–978.

31. Globyte, V. and Joo, C. (2019) Single-molecule FRET studies of Cas9 endonuclease. Methods Enzymol, 616, 313–335.

32. Siddiqui-Jain, A., Grand, C.L., Bearss, D.J. and Hurley, L.H. (2002) Direct evidence for a G-quadruplex in a promoter region and its targeting with a small molecule to repress c-MYC transcription. Proceedings of the National Academy of Sciences, 99, 11593–11598.

33. Cogoi, S. and Xodo, L.E. (2006) G-quadruplex formaon within the promoter of the KRAS proto-oncogene and its effect on transcripon. Nucleic Acids Res, 34, 2536–2549.

34. Beals, N., Farhath, M.M., Kharel, P., Croos, B., Mahendran, T., Johnson, J. and Basu, S. (2021) Rationally designed DNA therapeutics can modulate human tyrosine hydroxylase expression by controlling specific G-quadruplex formaon in its promoter. Mol Ther, 10.1016/j.ymthe.2021.05.013.

35. Farhath, M.M., Thompson, M., Ray, S., Sewell, A., Balci, H. and Basu, S. (2015) G-Quadruplex-Enabling Sequence within the Human Tyrosine Hydroxylase Promoter Differentially Regulates Transcription. Biochemistry, 54, 5533–5545.

36. Siddiqui-Jain, A., Grand, C.L., Bearss, D.J. and Hurley, L.H. (2002) Direct evidence for a G-quadruplex in a promoter region and its targeting with a small molecule to repress c-MYC transcription. Proc Natl Acad Sci U S A, 99, 11593–11598.

37. Furlong, R.A., Rubinsztein, J.S., Ho, L., Walsh, C., Coleman, T.A., Muir, W.J., Paykel, E.S., Blackwood, D.H. and Rubinsztein, D.C. (1999) Analysis and metaanalysis of two polymorphisms within the tyrosine hydroxylase gene in bipolar and unipolar affective disorders. Am J Med Genet, 88, 88–94.

38. Ishiguro, H., Arinami, T., Saito, T., Akazawa, S., Enomoto, M., Mitushio, H., Fujishiro, H., Tada, K., Akimoto, Y., Mifune, H., et al. (1998) Systematic search for variations in the tyrosine hydroxylase gene and their associations with schizophrenia, affective disorders, and alcoholism. Am J Med Genet, 81, 388–396.

39. Kunugi, H., Kawada, Y., Hattori, M., Ueki, A., Otsuka, M. and Nanko, S. (1998) Association study of structural mutations of the tyrosine hydroxylase gene with schizophrenia and Parkinson’s disease. Am J Med Genet, 81, 131–133.

40. Marcu, K.B., Bossone, S.A. and Patel, A.J. (2003) myc FUNCTION AND REGULATION. 10.1146/annurev.bi.61.070192.004113, 61, 809–858.

41. Pelengaris, S., Rudolph, B. and Littlewood, T. (2000) Action of Myc in vivo — proliferation and apoptosis. Curr Opin Genet Dev, 10, 100–105.

42. Thompson, E.B. (1998) The many roles of c-myc in apoptosis. Annu Rev Physiol, 60, 575–600.

43. Gabay, M., Li, Y. and Felsher, D.W. (2014) MYC acvation is a hallmark of cancer initiation and maintenance. Cold Spring Harb Perspect Med, 4.

44. Dhanasekaran, R., Deutzmann, A., Mahauad-Fernandez, W.D., Hansen, A.S., Gouw, A.M. and Felsher, D.W. (2022) The MYC oncogene - the grand orchestrator of cancer growth and immune evasion. Nat Rev Clin Oncol, 19, 23–36.

45. Postel, E.H., Mango, S.E. and Flint, S.J. (1989) A Nuclease-Hypersensitive Element of the Human c-myc Promoter Interacts with a Transcription Initiation Factor. Mol Cell Biol, 9, 5123–5133.

46. Sakatsume, O., Tsutsui, H., Wang, Y., Gao, H., Tang, X., Yamauchi, T., Murata, T., Itakura, K. and Yokoyama, K.K. (1996) Binding of THZif-1, a MAZ-like Zinc Finger Protein to the Nuclease-hypersensitive Element in the Promoter Region of the c-MYC Protooncogene*. J Biol Chem, 271, 31322–31333.

47. Balci, H., Globyte, V. and Joo, C. (2021) Targeting G-quadruplex Forming Sequences with Cas9. ACS Chem Biol, 16, 596–603.

48. Maleki, P., Budhathoki, J.B., Roy, W.A. and Balci, H. (2017) A practical guide to studying G-quadruplex structures using single-molecule FRET. Molecular Genetics and Genomics, 292, 483–498.

49. Broxson, C., Beckett, J. and Tornaletti, S. (2011) Transcription arrest by a G quadruplex forming-trinucleotide repeat sequence from the human c-myb gene. Biochemistry, 50, 4162–4172.

50. Bhattacharyya, D., Mirihana Arachchilage, G. and Basu, S. (2016) Metal Cations in G-Quadruplex Folding and Stability. Front Chem, 4, 38.

51. Vigouroux, A., Oldewurtel, E., Cui, L., Bikard, D. and van Teeffelen, S. (2018) Tuning dCas9’s ability to block transcription enables robust, noiseless knockdown of bacterial genes. Mol Syst Biol, 14, e7899.

52. Clarke, R., Heler, R., MacDougall, M.S., Yeo, N.C., Chavez, A., Regan, M., Hanakahi, L., Church, G.M., Marraffini, L.A. and Merrill, B.J. (2018) Enhanced Bacterial Immunity and Mammalian Genome Editing via RNA-Polymerase-Mediated Dislodging of Cas9 from Double-Strand DNA Breaks. Mol Cell, 71, 42–55.e8.

53. Hall, P.M., Inman, J.T., Fulbright, R.M., Le, T.T., Brewer, J.J., Lambert, G., Darst, S.A. and Wang, M.D. (2022) Polarity of the CRISPR roadblock to transcription. Nature Structural & Molecular Biology |, 29, 1217–1227.

