## Supplementary Information for "Stalling of Transcription by Putative G-quadruplex Sequences and CRISPR-dCas9"

**Table S1** *Oligonucleotide sequences used for single molecule FRET assay (Figure 3 of manuscript) for Tyrosine Hydroxylase system.*

| Name | Sequence (5' to 3') |
|---|---|
| **Non-Template Strand (C-rich)** | GCTAATACGA CTCACTATAG GAAAG CCCCCTCTGGGTCCCCCACCT TCCCCTCC TTACATCCCCCACCCCTGCCTGCTG |
| **Template Strand (G-rich)** | CAGCAGGCAGGGGTGGGGGATGTAAGGAGGGGAAGGTGGGGGACCCAGAGGG GGCTTTCCTATAGTGAGTCGTATTAGCTTTGGCGACGGCAGCGAGGC |
| **Top18-Biotin:** | Biotin- GCC TCG CTG CCG TCG CCA |

**T**: Cy5 Labeled
**A**: Cy3 Labeled

The orange segments are complementary to each other and form an 18 bp duplex that is used to attach the DNA construct to the surface.

The blue segments are complementary to each other and form the promoter site for T7 RNA polymerase.

The underlined G-rich sequence in the Template Strand is the putative quadruplex sequence and the underlined C-rich sequence in the Non-Template Strand is its complementary sequence.

**Table S2** *Oligonucleotide sequences used for in vitro transcription assay for Tyrosine Hydroxylase (constructs of Figure 4-7 of the manuscript).*

| Name | Sequence (5' to 3') |
|---|---|
| **200 bp Construct: Template Strand (Fig. 5)** | GTTATGCCTCCATCAGGCACAGCAGGCAGGGGTGGGGGATGTAAGGAGGGGAA GGTGGGGGACCCAGAGGGGGCTTTGACGTCAGCTCAGCTTATAAGAGGCTGCTG GGTATTCTCCATTATGATAGCTATTTAACTCGACTAAATCTCATGATACCTATGT AACTCGACTAAATGTTCCCTATAGTGAGTCGTATTAGC |
| **200 bp Construct: Non-Template Strand (Fig. 5)** | GCTAATACGACTCACTATAGGGAACATTTAGTCGAGTTACATAGGTATCATGAG ATTTAGTCGAGTTAAATAGCTATCATAATGGAGAATACCCAGCAGCCTCTTATA AGCTGAGCTGACGTCAAAGCCCCCTCTGGGTCCCCCACCTTCCCCTCCTTACATC CCCCACCCCTGCCTGCTGTGCCTGATGGAGGCATAAC |
| **135 bp Construct: Template Strand (Fig. 6)** | GCTGAGCTGACGTCAAAGCCCCCTCTGGGTCCCCCACCTTCCCCTCCTTACATCC CCCACCCCTGCCTGCTGTGCCTGATGGAGGCGGGGGTGGGTGAGGAAGGAGGG AGGCGCCCTATAGTGAGTCGTATTAGC |
| **135 bp Construct: Non-Template Strand (Fig. 6)** | GCTAATACGACTCACTATAGGGCGCCTCCCTCCTTCCTCACCCACCCCCGCCTCC ATCAGGCACAGCAGGCAGGGGTGGGGGATGTAAGGAGGGGAAGGTGGGGGAC CCAGAGGGGGCTTTGACGTCAGCTCAGC |
| **125 bp Construct: Template Strand (Fig. 4)** | CCTCCATCAGGCACAGCAGGCAGGGGTGGGGGATGTAAGGAGGGGAAGGTGGG GGACCCAGAGGGGGCTTTGACGTCAGCTCAGCTTATAAGAGGCTGCTGGGCCCT ATAGTGAGTCGTATTAGC |
| **125 bp Construct: Non-Template Strand (Fig. 4)** | GCTAATACGACTCACTATAGGGCCCAGCAGCCTCTTATAAGCTGAGCTGACGTC AAAGCCCCCTCTGGGTCCCCCACCTTCCCCTCCTTACATCCCCCACCCCTGCCTG CTGTGCCTGATGGAGG |
| **TracrRNA** | GGAACCAUUCAAAACAGCAUAGCAAGUUAAAAUAAGGCUAGUCCGUUAUCAA CUUGAAAAAGUGGCACCGAGUCGGUGCUUUUUUU |
| **cR-1** | N6-GGCCCCUGCCUGCUGUGCCUGAGUUUUAGAGCUAUGCUGUUUUG |
| **cR-2** | N6-GGUCAGGCACAGCAGGCAGGGGGUUUUAGAGCUAUGCUGUUUUG |
| **cR-3** | GGGCUGACGUCAAAGCCCCCUCGUUUUAGAGCUAUGCUGUUUUG |
| **cR-4** | N6-GGGAAGGUGGGGGACCCAGAGGGUUUUAGAGCUAUGCUGUUUUG |

**Table S3** *Oligonucleotide sequences used for in vitro transcription assay for c-Myc system (construct of Figure 8 of manuscript).*

| Name | Sequence (5' to 3') |
|---|---|
| **Non-Template Strand (C-rich)** | GCTAATACGACTCACTATAGGAGCAAAAGAAAATGGTAGGCGCGCGTAGTTAATTCATGCGGCTCTCTTACTCTGTTTACATCCTAGAGCTAGAGTGCTCGGCTGCCCGGCTGAGTCTCCTCCCCACCTTCCCCACCCTCCCCACCCTCCCCATAAGCGCCCCTCCCGGGTTCCCAAAGCAGAGGGCGTGGGGG |
| **Template Strand (G-rich)** | CCCCCACGCCCTCTGCTTTGGGAACCCGGGAGGGGCGCTTATGGGGAGGGTGGGGAGGGTGGGGAAGGTGGGGAGGAGACTCAGCCGGGCAGCCGAGCACTCTAGCTCTAGGATGTAAACAGAGTAAGAGAGCCGCATGAATTAACTACGCGCGCCTACCATTTTCTTTTGCTCCTATAGTGAGTCGTATTAGC |
| **TracrRNA** | GGAACCAUUCAAAACAGCAUAGCAAGUUAAAAUAAGGCUAGUCCGUUAUCAACUUGAAAAAGUGGCACCGAGUCGGUGCUUUUUUU |
| **cR-1** | GGCTCCCCATAAGCGCCCCTCCGUUUUAGAGCUAUGCUGUUUUG |
| **cR-2** | GGAGGGTGGGGAAGGTGGGGGUUUUAGAGCUAUGCUGUUUUG |
| **cR-3** | GGACCCGGGAGGGGCGCTTATGGUUUAGAGCUAUGCUGUUUG |
| **cR-4** | GGCAGCCGAGCACTCTAGCTCTGUUUUAGAGCUAUGCUGUUUUG |
| **cR-5** | GGCGCCCTCTGCTTTGGGAACCGUUUUAGAGCUAUGCUGUUUUG |
| **cR-6** | GGAGCTAGAGTGCTCGGCTGCCGUUUUAGAGCUAUGCUGUUUUG |
| **cR-7** | GGTCCCGGGTTCCCAAAGCAGAGUUUUAGAGCUAUGCUGUUUUG |

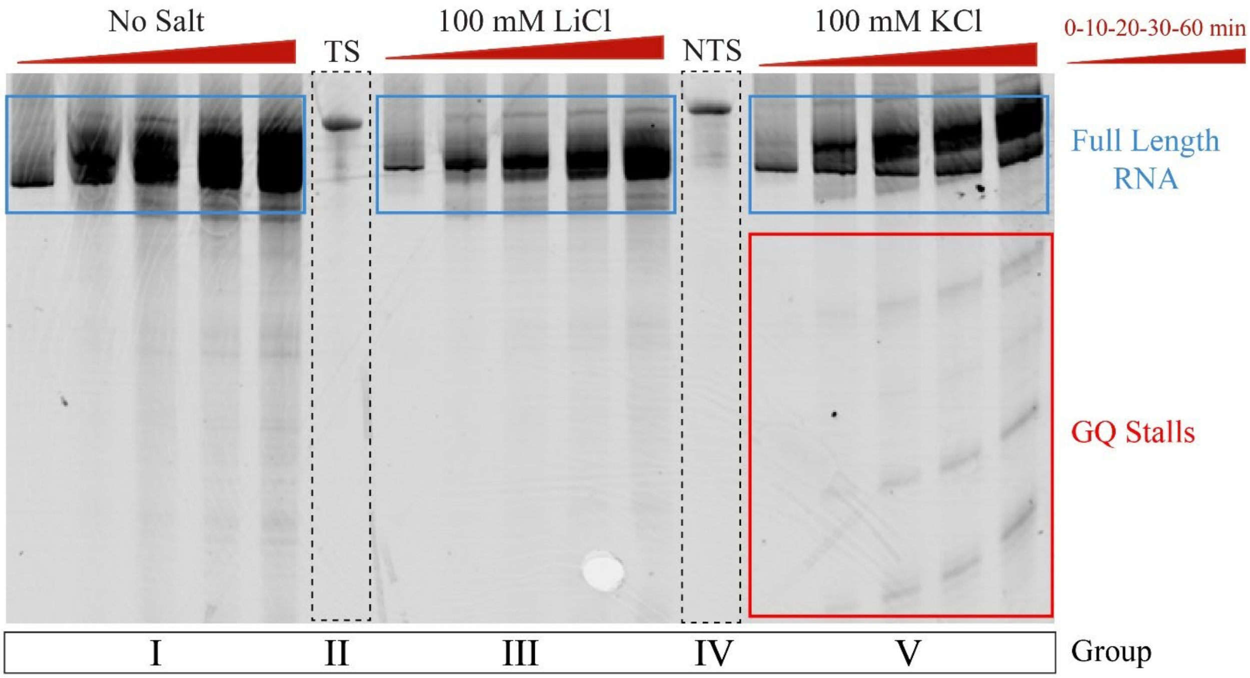


**Figure S1.** Denaturing PAGE images showing truncated RNA products due to GQ stalls. These stalls are prominent in KCl but much weaker in LiCl and no salt conditions. The lane indicated with TS is the C-rich template strand and NTS is the G-rich non-template strand.

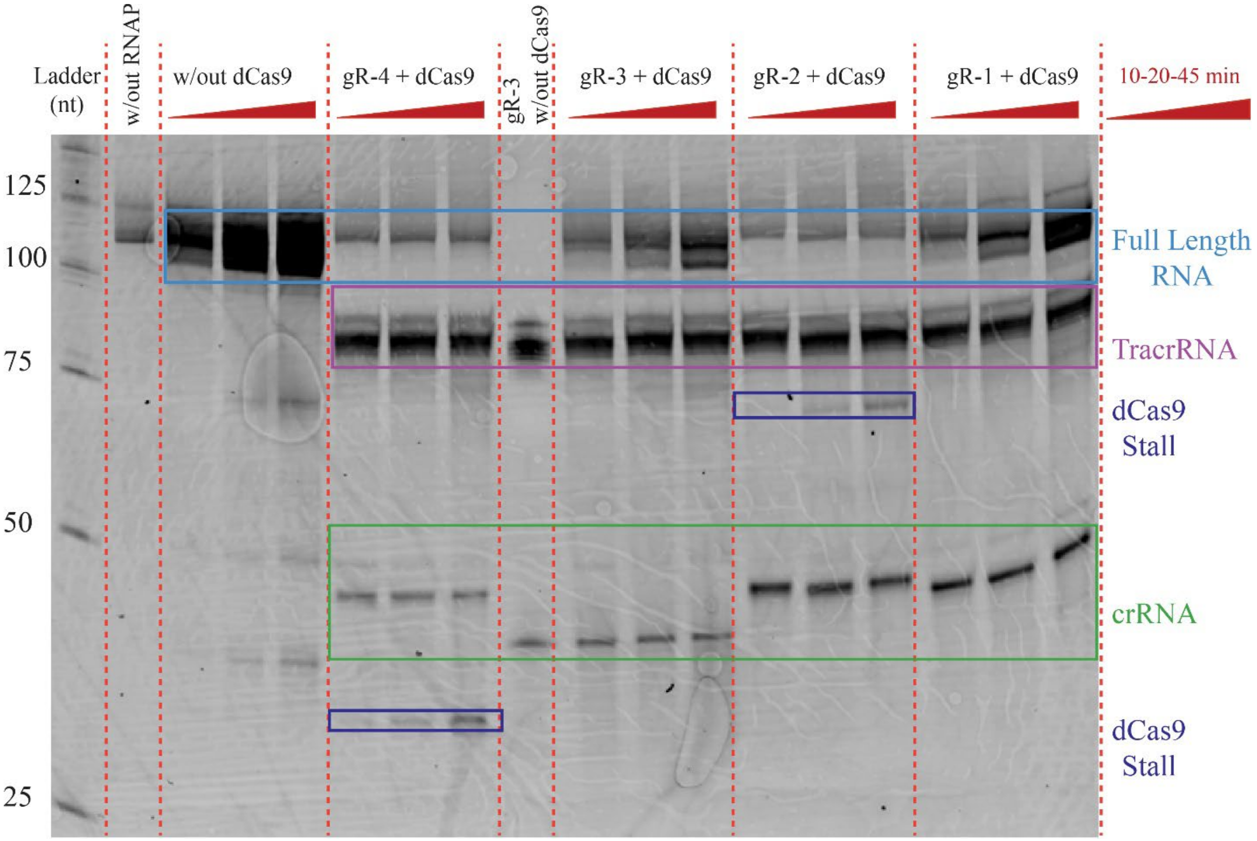


**Figure S2.** Original (uncut) version of the gel image shown in Figure 4 (PQS in the template strand for the tyrosine hydroxylase system).

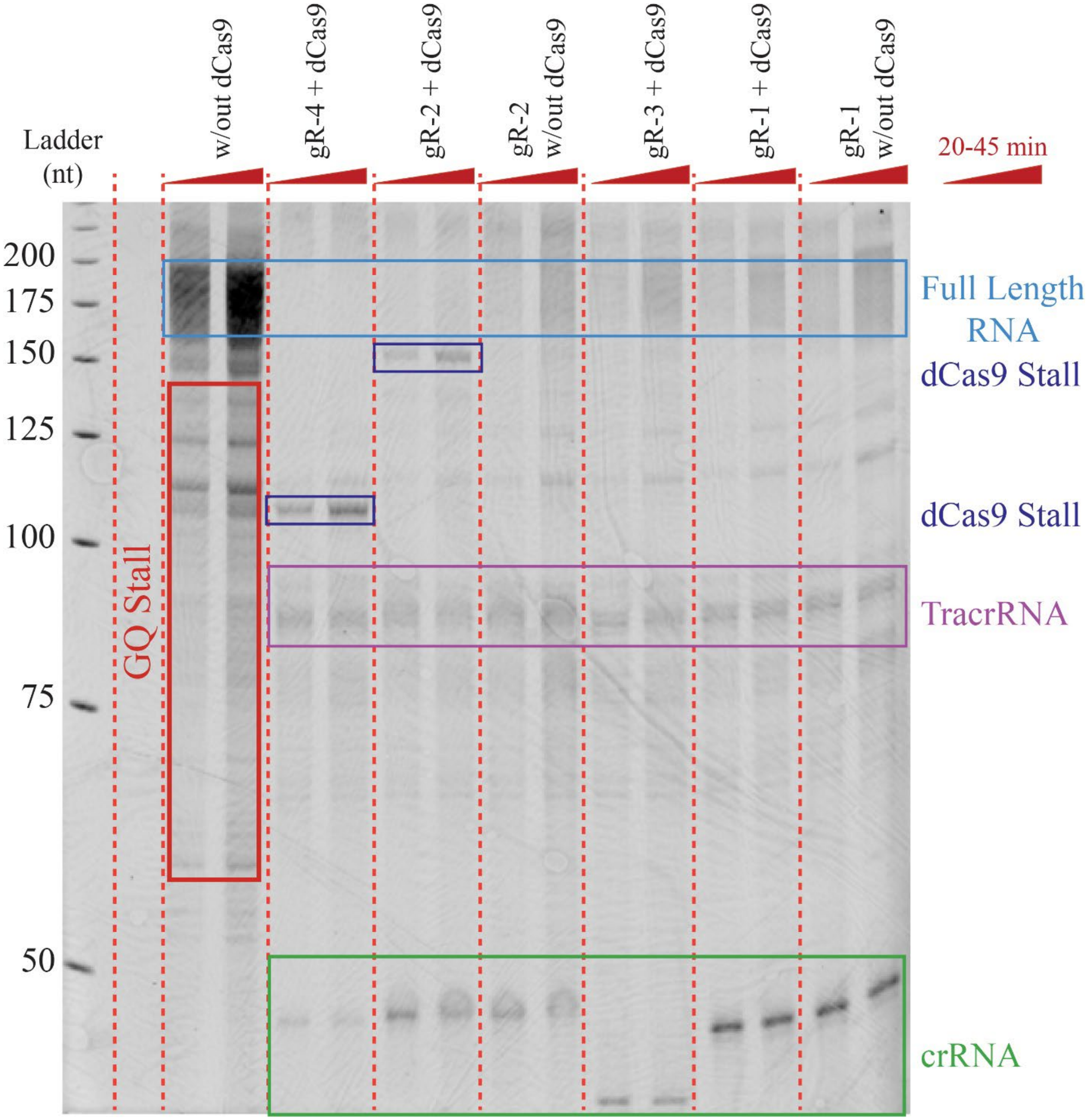


**Figure S3.** Original (uncut) version of the gel image shown in Figure 5 (PQS in the template strand for 200 bp long DNA construct for the tyrosine hydroxylase system).

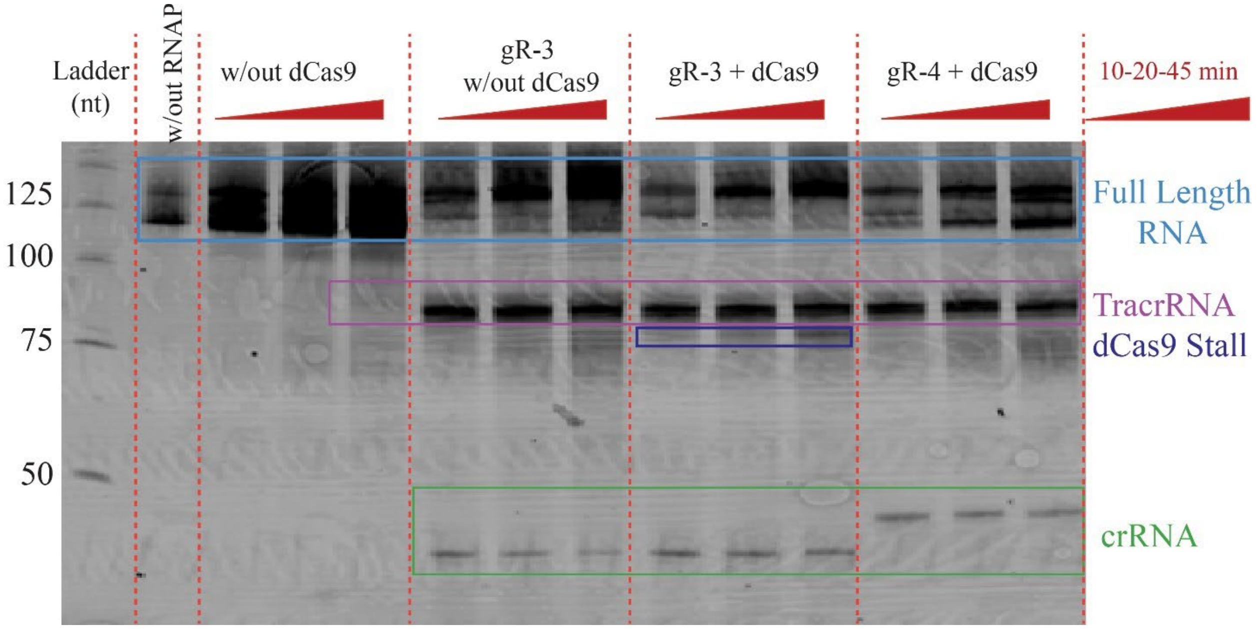


**Figure S4.** Original (uncut) version of the gel image shown in Figure 6 (PQS in the non-template strand for the tyrosine hydroxylase system).

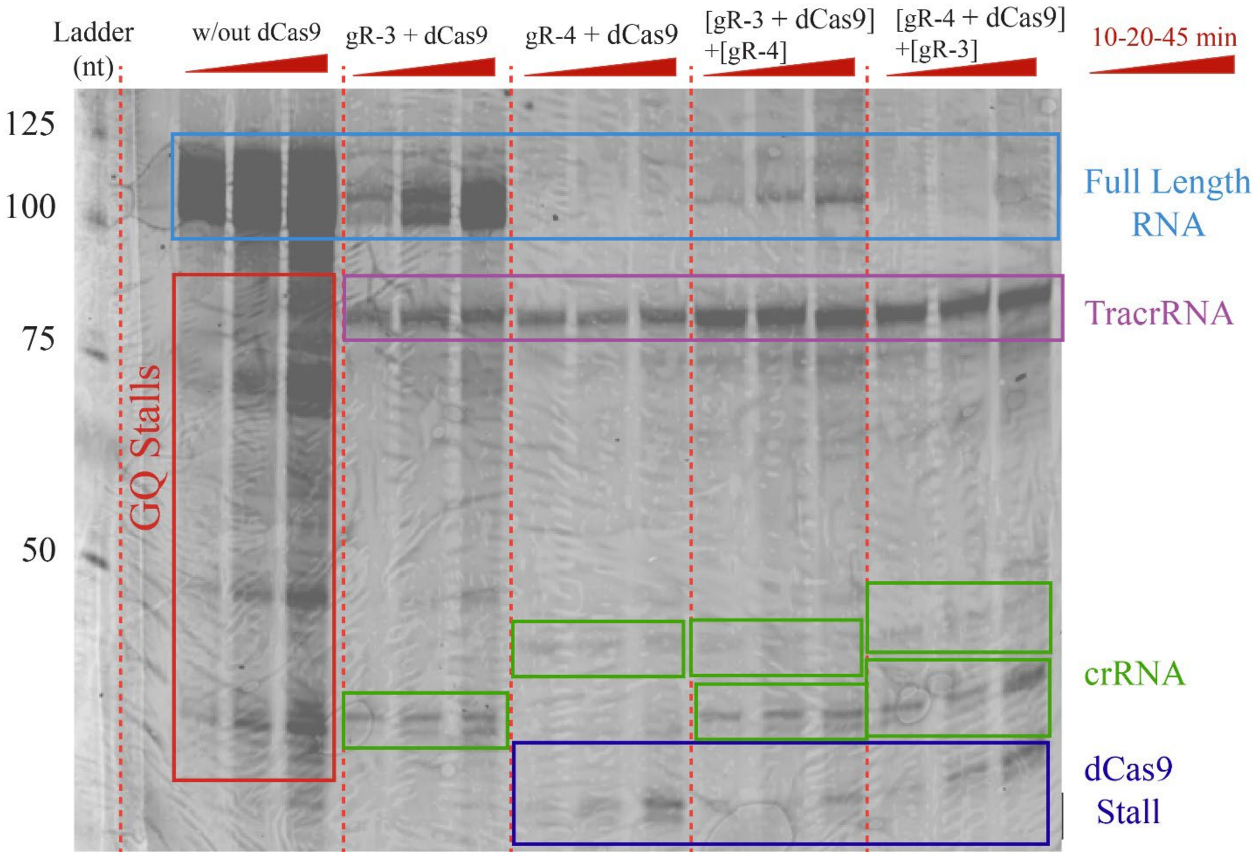


**Figure S5.** Original (uncut) version of the gel image shown in Figure 7 (PQS in the template strand for the tyrosine hydroxylase system, competition assay).

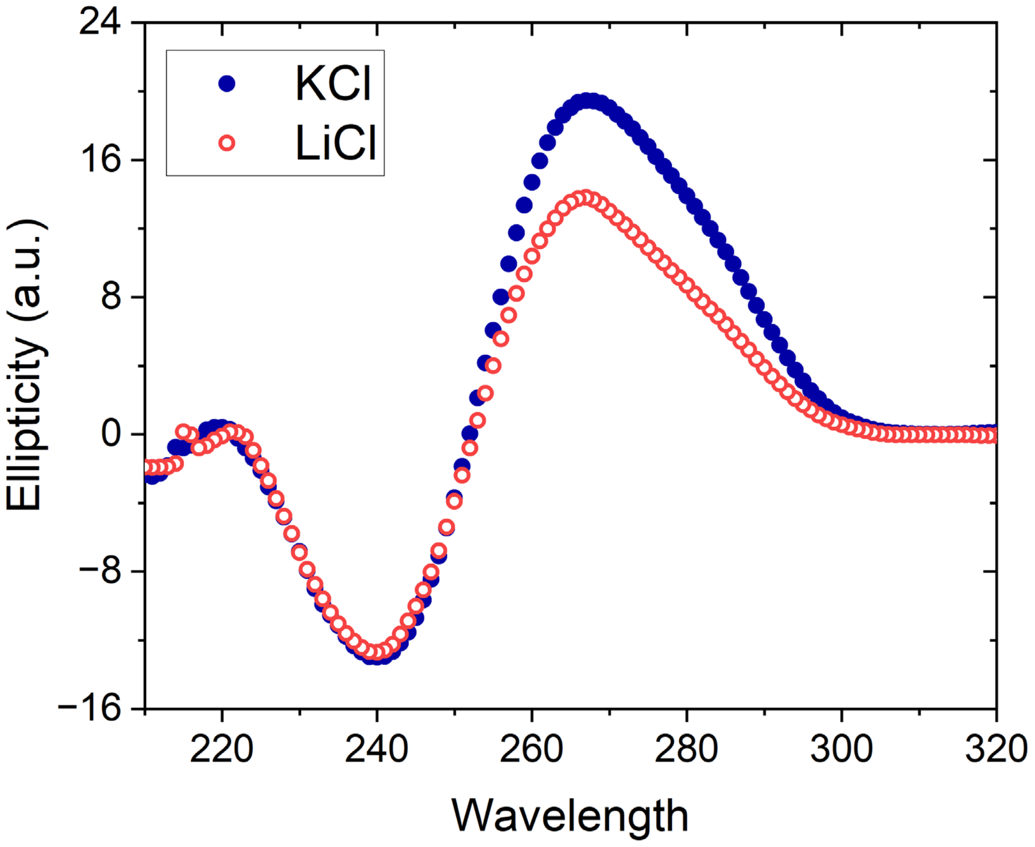


**Figure S6.** Circular dichroism spectrum of the c-Myc DNA construct in 100 mM KCl or 100 mM LiCl. The peak at 265 nm and trough at 240 nm are consistent with parallel conformation. The more prominent peak in KCl suggests a more prominent GQ formation in KCl compared to LiCl.
